# Involvement of neurons in the non-human primate anterior striatum in proactive inhibition

**DOI:** 10.1101/2024.04.24.591009

**Authors:** Atsushi Yoshida, Okihide Hikosaka

## Abstract

Behaving as desired requires selecting the appropriate behavior and inhibiting the selection of inappropriate behavior. This inhibitory function involves multiple processes, such as reactive and proactive inhibition, instead of a single process. In this study, macaque monkeys were required to perform a task in which they had to sequentially select (accept) or refuse (reject) a choice. Neural activity was recorded from the anterior striatum, which is considered to be involved in behavioral inhibition, focusing on the distinction between proactive and reactive inhibitions. We identified neurons with significant activity changes during the rejection of bad objects. Cluster analysis revealed three distinct groups, of which one showed obviously increased activity during object rejection, suggesting its involvement in proactive inhibition. This activity pattern was consistent irrespective of the rejection method, indicating a role beyond mere saccadic suppression. Furthermore, minimal activity changes during the fixation task indicated that these neurons were not primarily involved in reactive inhibition. In conclusion, these findings suggest that the anterior striatum plays a crucial role in cognitive control and orchestrates goal-directed behavior through proactive inhibition, which may be critical in understanding the mechanisms of behavioral inhibition dysfunction that occur in patients with basal ganglia disease.

## Introduction

Achieving goals requires selecting the desired action and suppressing other compelling actions. The inhibition of actions is multifaceted, and there are distinctions between automatic and volitional suppressions, as well as reactive and proactive suppressions (Sumner et al., 2007; Boy et al., 2010; Aron, 2011; Verbruggen & Logan, 2014; Zandbelt & Vink, 2010; Jahanshahi et al., 2015). Reactive inhibition is often triggered by external stimuli and is habitual and reflexive. For instance, a red traffic light prompts us to stop walking or apply the brakes while driving. In contrast, proactive inhibition is volitional and goal-directed, such as when students resist the temptation to play to focus on their studies. Understanding the specific brain regions that control these inhibitory functions is crucial to advancing our knowledge of cognitive control mechanisms.

Previous functional brain imaging studies in humans and physiological studies in experimental animals, including rodents and non-human primates, have implicated an indirect pathway in the basal ganglia for these inhibitory functions. This pathway includes the striatum (caudate nucleus and putamen), globus pallidus external segment (GPe), subthalamic nucleus (STN), and globus pallidus internal segment or substantia nigra pars reticulata (SNr). Specifically, the striatum and the STN have been associated with reactive inhibition (Aron & Poldrack, 2006; Aron & Poldrack, 2007; Li et al., 2008), and the striatum has been suggested to play a crucial role in proactive suppression (Vink et al., 2005; Jahfari et al., 2012; Chikazoe et al., 2009; Smittenaar et al., 2013; Majid et al., 2013; Schel et al., 2014). Neurophysiological studies involving non-human primates initially demonstrated the involvement of the striatum in reactive inhibition (Apicella et al., 1992; Hori et al., 2009), while subsequent studies have indicated the involvement of both the striatum and GPe in proactive inhibition (Watanabe & Munoz, 2010; Yoshida & Tanaka, 2009; Yoshida & Tanaka, 2016). However, the anti-saccade task used in these studies, which requires both the facilitation of desired choices and the inhibition of unwanted behaviors, makes it challenging to discern specific neural representations.

To address this issue, we developed a choice task with sequential presentation of alternatives, allowing for the separation of choice and rejection actions. By recording neuronal activity from the GPe from macaque monkeys while performing this task, we found a subset of GPe neurons significantly changed their neuronal activity when rejecting an object choice (Yoshida & Hikosaka, 2023). Furthermore, by combining it with a fixation task that required inhibition of the reflexive saccades (reactive inhibition), we showed that this group of neurons in the GPe may be involved in proactive inhibition.

Our previous study (Yoshida and Hikosaka, 2023) found most neurons involved in proactive inhibition in the dorsal part of the GPe; thus, we recorded neural activity mainly in the anterior and body parts of the striatum, which were thought to project to this area. The purpose of this study was to investigate whether the neurons in the striatum were involved in proactive inhibition using the choice and fixation tasks used in our previous study. Given that the firing rates of the GPe neurons decreased during proactive inhibition, we focused our neural activity recordings on the medium spiny neurons in the striatum that sent inhibitory projections to the GPe (Precht and Yoshida, 1971).

## Materials and Methods

### Animal preparation

All animal care and experimental procedures were approved by the National Eye Institute Animal Care and Use Committee and complied with the Public Health Service Policy on Human Care and Use of Laboratory Animals.

Two male macaque monkeys (Macaca mulatta, aged 9 years, 9–10 kg body weight), referred to as Monkeys C and S, were used. After acclimation to the chair, a plastic head holder and recording chambers were securely affixed to the craniums of both subjects with ceramic screws and dental acrylic under isoflurane anesthesia administered under sterile conditions. Pain relief was provided with analgesics for five days during and after the surgical procedure. Training for oculomotor tasks commenced after surgical recovery. The daily water intake of the monkeys was controlled to motivate them to perform behavioral tasks. The monkey heads were immobilized using a primate chair throughout the training and experiments. An eye-tracking device (EyeLink 1000; SR Research, Ontario, Canada) recorded the eye movements at a frequency of 1000 Hz.

### Experimental design

The trials were conducted in a secluded, light-controlled, and soundproof environment. The participants were positioned on a restraint device facing an interactive display. Visual stimuli were projected onto the screen in front of the participants using an active matrix liquid crystal display projector (PJ658; ViewSonic Corp., Brea, CA, USA). A custom-made C++-based experimental data acquisition system (Blip; available at http://www.robili.sblip/) was used to control the behavioral tasks. This study employed two distinct behavioral tasks: choice and fixation.

### Choice and fixation tasks

We used the choice and fixation tasks. In the choice task, circular greyscale aerial images (radius: 25°) from OpenAerialMap, referred to as scenes, were presented as the background on the screen (Figure 1A). Computer-generated multi-colored fractal objects were utilized as targets (radius: 5°) (Yamamoto et al., 2013). One of the six sets of scenes was randomly chosen during each single-unit recording session (Figure 1B). Each set contained four different scenes featuring one “good” fractal object (associated with reward delivery) and one “bad” object (not associated with a reward). The value rationings to the stimuli remained unchanged in the “stable” scenes (1 and 2) but were changeable in the “flexible” scenes (3 and 4). This design aimed to discern the relationship of these neuronal responses to the features of the objects or their value assignments.

**Figure 1.**
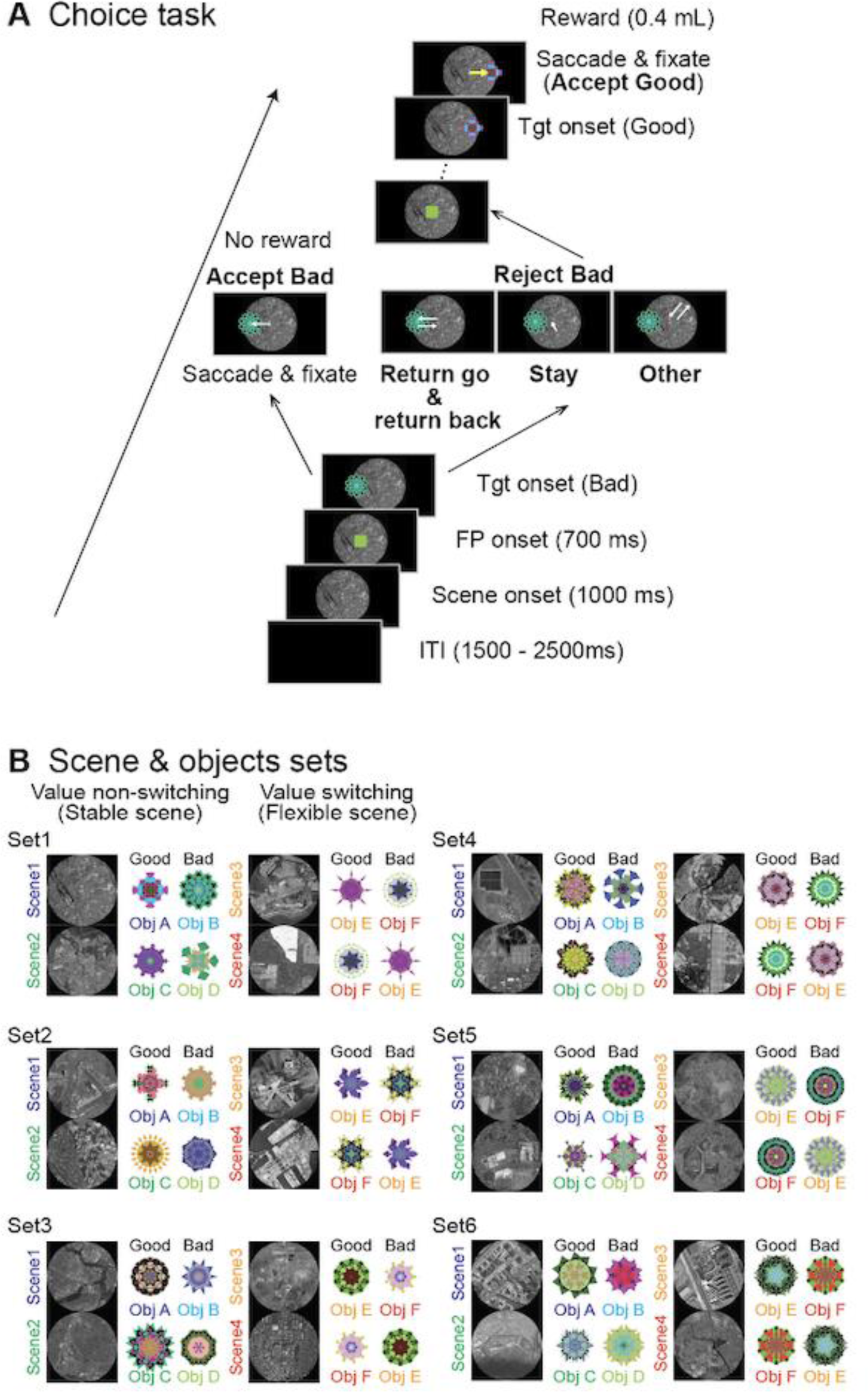
Task procedures. (**A**) Procedure for the choice task. The monkeys are considered to have chosen a fractal object presented to them if they make a saccade toward it and fixated on it for more than 400 ms (*Accept*). They reject the object (*Reject*) by making a return saccade (*Return*), looking around the center point (*Stay*), or making a saccade away from the bad object (*Other*). **(B)** All six sets used for recording from a single neuron. For neuronal recording from an individual neuron, only one set is used. One set includes four *scenes*, and each *scene* has one good (rewarded) and one bad (non-rewarded) fractal object during the choice task. In *scenes* 1 and 2, the object values are always the same (stable), while the same two objects are used in *scenes* 3 and 4, but their values are replaced (flexible) in each *scene*.

The task sequence started with the unveiling of a scene following a preparatory period (1500–2000 ms). The subject saw the scene freely before a fixation point (FP) emerged at the center of the display (1000 ms). After maintaining the stare at the FP (700 ms), one of the fractal objects, either good or bad, randomly and sequentially appeared as a target after the FP offset. The target was randomly shown in one of six positions (eccentricity: 15°; angles: 0°, 45°, 135°, 180°, 225°, and 315°). If the participant made a saccade toward the target and fixated on it for more than 400 ms, this action was recorded as object acceptance (accept), resulting in the delivery of a reward (0.4 mL) for a good object. No reward was given for bad objects. If the subject kept looking around the center point (stay), made a saccade to the target but returned their gaze to the center point within 400 ms (return), or looked away from the target (other), the FP would reappear. These actions were considered when the participants refused to select an object (rejected).

The range of the target was set to a square (8° per side). If the eye position of the subject went outside the target range for less than 400 ms after making a saccade to the target, the target was gone and an FP appeared again. As a result, when the monkeys rejected bad objects by staying or other means, the waiting time for the next FP to be presented was 1000 ms, whereas the waiting time was less than 1000 ms if they rejected bad objects by returning. When the subject rejected bad objects by returning, two saccades occurred: one was toward the object (return go), and the other was from the object to the center (return back). Another target was presented when the subjects were gazing at the FP. This procedure was repeated 15 times until the participants selected the target object.

The aim of the fixation task was to elucidate whether the neuronal activity was related to suppressing saccades to presented objects. This task followed the choice task in which individual neurons were isolated. In this task, both fractal objects (one good and one bad object) from scene 1 of the choice task were displayed alternately 2–4 times to the left or right of the FP; each was displayed for 400 ms with a 400-ms interstimulus interval. The subjects were required to keep their gaze on a white fixation point (FP) during the simultaneous display of the fractal object. A reward (0.4 mL) was given 300 ms after the presentation of the last object. Saccades toward the object stimuli were considered fixation break errors.

### MRI

After the surgery to implant recording chambers, magnetic resonance imaging (MRI) was performed to delineate cerebral structures and apertures in the grid with a gadolinium-enhanced contrast medium (Magnevist, Bayer Healthcare Pharmaceuticals, Wayne, NJ, USA) introduced into the chambers. The recording sites were pinpointed and reconstructed from the MRI data using a high-definition 3T MR apparatus (MAGNETOM Prisma; Siemens Healthcare, Erlangen, Germany). This process involved the acquisition of both three-dimensional T1-weighted (T1w, MPRAGE) and T2-weighted (T2w, SPACE) sequences, each with a uniform voxel dimension (0.5 mm) (Mugler et al., 2000).

To illustrate the locations of the recorded task-related neurons in the striatum, we segmented the region of the striatum automatically using a pipeline with the AFNI software (Cox, 1996; Jung et al., 2021; Seidlitz et al., 2018). Each monkey T1w image was aligned to a standard NIMH macaque templates (NMT v2.0) using the AFNI macaque pipeline. Then, the subcortical atlas of the rhesus macaque (SARM) for NMT (Hartig et al., 2021) was inversely transformed into a native T1w image. We compared the segmented images of the striatum with the quantitative susceptibility mapping images of the subjects to confirm the validity of the former; the former can visualize the structure of the striatum more clearly than T1w and T2w images (Yoshida et al., 2021; Dadarwal et al., 2022). We used the 3D Slicer software (version 5.2.2) (Fedorov et al., 2012) and Blender software (version 3.5) to visualize the locations of the recorded task-related neurons.

### Neuronal recording procedure

Neuronal activity was captured via single-unit recordings with epoxy-or glass-coated tungsten electrodes with impedances between 1 MΩ and 9 MΩ (Frederick Haer & Co., Bowdoinham, ME, USA; Alpha Omega Engineering, Nazareth, Israel). The electrodes were carefully positioned within the cerebral tissue using a precision oil-driven micromanipulator (model MO-973A; Narishige, Tokyo, Japan) that maneuvered the electrode through a guide tube made of 23-gauge stainless steel. The procured neural signals were amplified and filtered (0.3–10 kHz; A-M Systems Inc., Carlsborg, WA, USA). Once we succeeded in the isolation of neuronal signals, the choice task was initiated. The neuronal recording was continued if neuronal activity was modulated at the target onset.

To focus on the medium spiny neurons in the striatum, only neurons with baseline firing rates of less than 10 spikes/s were included in this study (Aosaki et al., 1995). The average baseline activity of neurons was calculated for 1000 ms between 1500 ms and 500 ms before the scene was presented. Thus, neurons with a high firing frequency exceeding 10 spikes/s were considered fast-spiking interneurons and excluded from the present study.

### Data analysis and Statistical analysis

All behavioral and neurophysiological data were preprocessed with MATLAB 2022b (MathWorks Inc., Natick, MA, USA).

#### Behavior

In the choice task, saccade onsets toward good and bad objects were defined as the eye speed exceeding 40°/s within 400 ms of the target onset. During the fixation task, saccades to the presented objects were detected offline. They were considered fixation break errors.

Welch’s t-tests were performed for each scene condition to compare the saccade reaction times for the good and bad objects (Figure 2A). During this analysis, only the reaction times of the saccades toward bad objects (return go) were included, and those for returning from the bad target (return back) were not included. To determine whether the proportion of monkeys choosing to stay for a bad object was higher for scenes 1 and 2, where the object values remained constant, Fisher’s exact test was conducted (Figure 2B).

**Figure 2.**
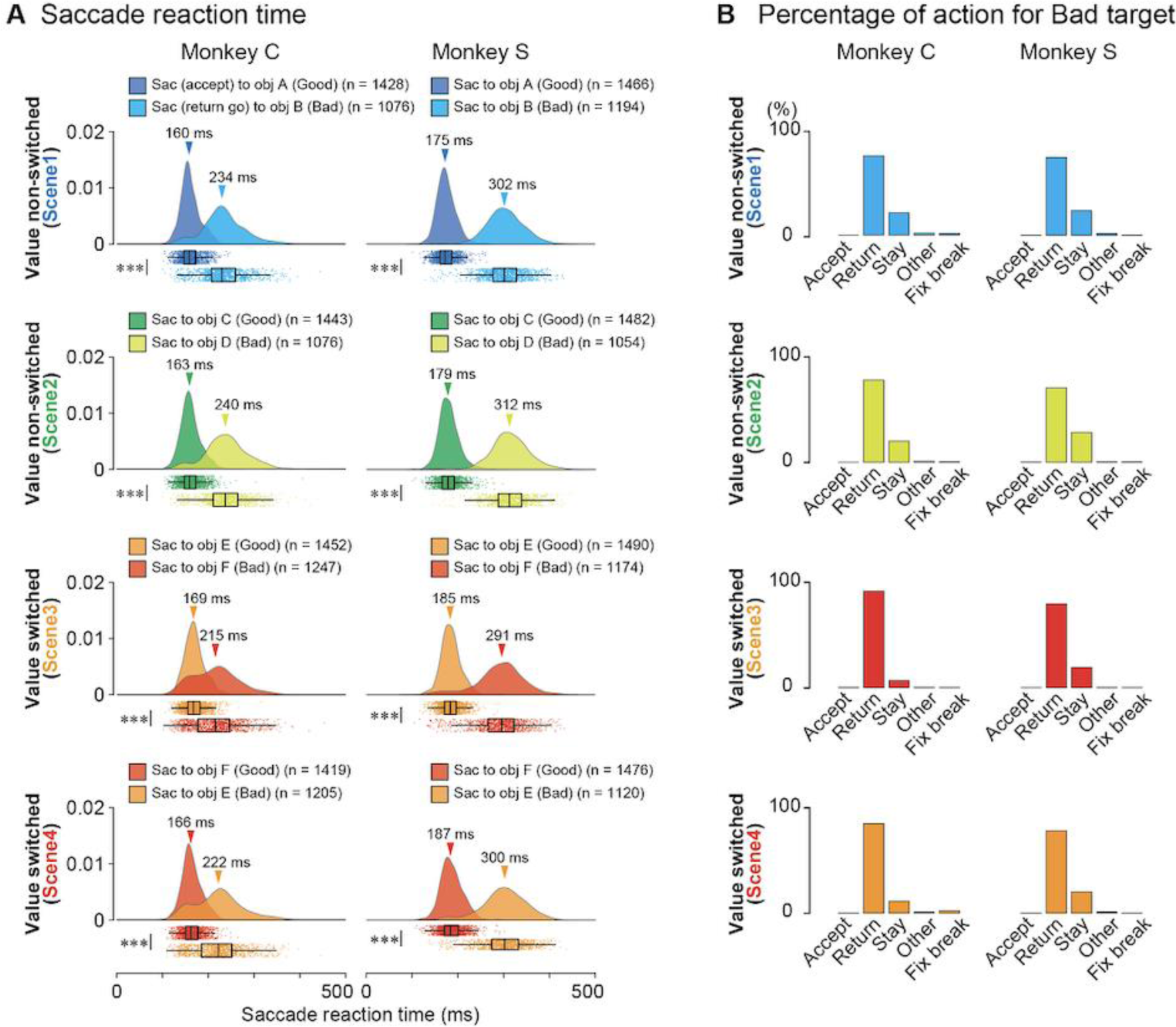
Behavioral results. **(A)** Raincloud plots of saccade reaction times for good or bad objects in *scenes* 1-4 during the choice task of the two monkeys, C and S. In each panel, the upper “cloud” illustrates the probability distribution of reaction times, and the lower “rain” shows the raw plots of individual reaction times. In all the *scenes,* the reaction times of the two monkeys toward good objects are significantly shorter than those toward bad objects (*p* = 0 for all *scenes* for both monkeys, Welch’s t-tests). **(B)** Proportions of selected actions, Accept, Return, Stay, Other, or fixation break error (Fix break) for bad objects in *scenes* 1-4 during the choice task. FP, fixation point; ITI, inter-trial interval; Obj, object; Tgt, target

#### Classification of striatal neurons

The sample size was not determined by statistical methods. Instead, the number of recorded neurons was determined based on a previous study (Watanabe & Munoz, 2010), which recorded the activities of striatal neurons of macaques while they performed an anti-saccade task. All statistical analyses were performed with the preprocessed data using the R software (version 4.2.2).

We employed the k-means clustering method to categorize neurons based on their activities when good or bad contralateral objects were presented. This process involved standardizing the activity of each neuron using a Z-transform. The average neuronal activity in response to the presentation of either a good or a bad object in the contralateral direction was computed. This calculation had a 200-ms window, extending from 100 ms to 300 ms after the onset of object presentation (Figure 3A). To determine the optimal number of clusters for k-means classification, a silhouette analysis was initially conducted by setting the number of clusters (K) to three. This preliminary step informed us of the number of clusters for our subsequent use of the k-means method for the final classification of neurons.

**Figure 3.**
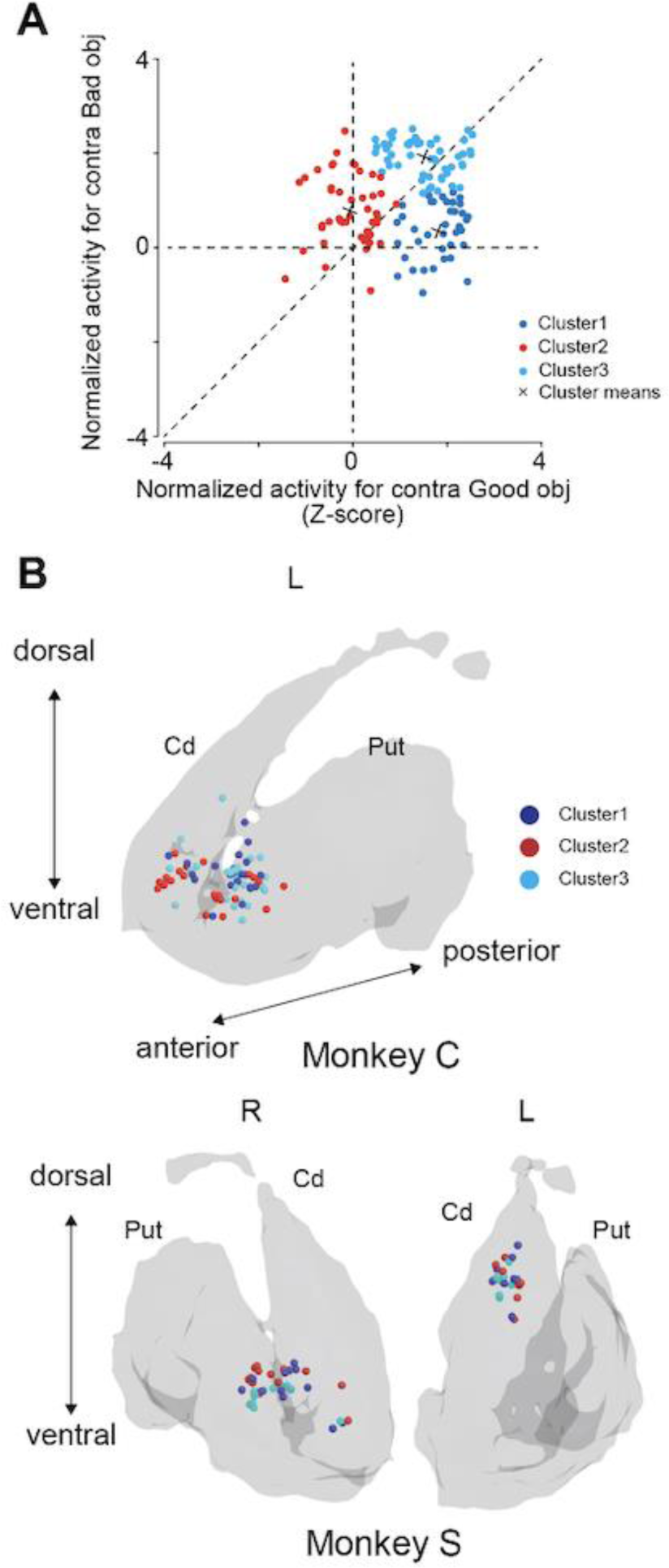
Scatter plots of clustering and recording sites in the striatum. **(A)** Two-dimensional scatter plots of the Z-transformed mean firing rate for individual neurons during 200 ms after 100 ms at the onsets of presentation of the good (X-axis) and bad (Y-axis) objects. Using these two-dimensional data, the neurons are divided into three groups using clustering methods. **(B)** Reconstruction of recording site of task-related striatal neurons of the two monkeys.

#### Comparisons of neuronal activity across conditions

The data from each neuron was aligned with the initiation of events (*scene*, target, and saccade onset). The time course of the neuronal response to event onset for each condition was investigated by calculating peristimulus time histograms (PSTHs) in 1-ms bins. It was smoothed with a spike density function using a Gaussian filter (σ = 20 ms). To visualize the neuronal activity of individual neurons on the color map, the Z-transformation of the activity in each neuron was performed to visualize the activities of individual neurons (Adler, et al., 2012; Kaplan et al., 2020). First, the firing rate of the baseline was calculated by averaging the firing rate within 500 ms before the *scene*, target, or saccade onset. Then, this baseline was subtracted from the smoothed PSTH and aligned with the initiation of events. Subsequently, a Z-score transformation was conducted for each PSTH; the baseline mean was subtracted from it, and the difference was divided by the standard deviation (SD).

To prevent false positives, generalized linear mixed-effects models (GLMMs) were used in this study to compare neuronal activities across conditions (Yu et al., 2022). First, we compared the full model containing explanatory variables as fixed effects for all statistical tests using the GLMM. All monkey, neuron, and experimental session IDs were included as random effects, which constituted the null model. A parametric bootstrap method was conducted to assess the goodness of fit of the models by performing 10,000 iterations and computing the *p*-value based on the difference in deviance between the two models. If the full model showed a significant fit, we conducted post hoc pairwise t-tests with Bonferroni corrections to further explore these differences. For GLMM, we used the *lme4* (Bates et al., 2014)*, pbkrtest* (Halekoh & Højsgaard, 2014), and *emmeans* (Lenth et al., 2019) packages in RStudio.

Neuronal activity at scene onset in the choice task was compared (Figures 5B, G, and K). A GLMM was employed to evaluate differences in neuronal activity at scene onset. The full and null models used for the comparison are as follows:

*Full Model: NeuronalActivity* ∼ *Scene* + (1|*monkey_ID*)

+ (1|*monkey_ID*: *Neuron_ID*),

*Null Model: NeuronalActivity* ∼ (1|*monkey_ID*) + (1|*monkey_ID*: *Neuron_ID*),

where *NeuronalActivity* is the mean PSTH of individual neurons during a 200 ms period, ranging from 100 ms to 300 ms after *scene* onset; *Scene* (1–4) is a fixed effect; and *monkey_ID* and *Neuron_ID* are random effects.

The level of statistical significance for the model comparison was set at α = 0.05.

Neuronal activities at target onset for the four scenes during the choice task were evaluated (Figures 5D, H, and L). A GLMM was conducted to examine the differences in neuronal activity at the target onset. The full and null models used for the comparison are as follows:

*Full Model: NeuronalActivity* ∼ *Scene* × *Value* × *Direction* + (1|*monkey_ID*)

+ (1|*monkey_ID*: *Neuron_ID*),

*Null Model: NeuronalActivity* ∼ (1|*monkey_ID*) + (1|*monkey_ID*: *Neuron_ID*),

where *NeuronalActivity* is the mean PSTH of individual neurons within 200 ms from 100 ms after the target onset; *Scene* (1–4), *Value* (good vs. bad), and *Direction* (contralateral vs. ipsilateral) are fixed effects, and *monkey_ID* and *Neuron_ID* are random effects.

Statistical significance for the model comparison was set at α = 0.05. We performed six pairwise t-tests to compare the mean neuronal activities for the good and bad objects during the choice task for each *scene* and six pairwise t-tests for comparison across *scenes*. Therefore, statistical significance for the post-hoc test was set at α = 0.05/12 with Bonferroni correction.

Neuronal activity at saccade onset during the choice task was assessed (Figures 6B, E, and H). Given that comparisons of the mean neuronal activity during the post-object onset period showed no apparent differences for the four scene conditions, the peri-saccade onset period was analyzed using averaged data from all *Scene* conditions. We employed a GLMM to investigate differences in neuronal activity during saccade onset. The full and null models used for the comparison are as follows:

*Full Model: NeuronalActivity* ∼ *Value* × *Direction* + (1|*monkey_ID*)

+ (1|*monkey_ID*: *Neuron_ID*),

*Null Model: NeuronalActivity* ∼ (1|*monkey_ID*) + (1|*monkey_ID*: *Neuron_ID*),

where *NeuronalActivity* refers to the mean PSTH of individual neurons during 200 ms beginning 100 ms after saccade onset; *Value* (good vs. bad) and *Direction* (contralateral vs. ipsilateral) are fixed effects, and *monkey_ID* and *Neuron_ID* are random effects. The level of statistical significance for the model comparison was set at α = 0.05. We performed six pairwise t-tests for comparison of mean neuronal activity for good and bad objects during the choice task, and statistical significance for the post-hoc test was set at α = 0.05/6 with Bonferroni correction.

For further analysis, a series of neuronal data analyses were conducted to ascertain the involvement of the recorded neuronal activity in facilitating saccades (Figure 7). Initially, the data recorded from each neuron were segmented into four groups based on the saccade reaction times in each trial under the four conditions (contralateral/ipsilateral versus good/bad object).

These groups were arranged in ascending order of saccade latency, with Group 1 having the shortest latency and the subsequent groups having progressively longer latencies. The average neuronal activity of each group was calculated. This calculation had a 200-ms window, from 100 ms to 300 ms following object presentation, and the resulting average neuronal activity was standardized using a Z-transformation. The correlation coefficients for individual neurons were computed to explore the relationship between saccade latency and neuronal activity. The coefficients were designed to assess the correlation between the group order (from Groups 1 to 4) and mean neuronal activity. Finally, Wilcoxon signed-rank tests were used to determine whether the median of the correlation coefficients differed significantly from zero across the datasets to establish the statistical significance of our findings.

Figures 8E, H, K, N, Q, and T show the comparison of neuronal activities at target onset while monkeys rejected bad objects by suppressing the saccade to a presented bad object (*stay*) or returning the gaze to the original center point without fixation for more than 400 ms after the saccade to the target (*return*) during the choice task and at object onset during the fixation task. By comparing the neural activities for these two cases, we determined whether the observed neural activity was related to the inhibition of the saccade or the rejection of the bad object. It was also determined whether the observed neural activity was related to proactive or reactive inhibition by comparing it with the neural activity during the fixation task. During the fixation task, the same objects were used (their values were constant during Scene 1 of the choice task). We compared the average neural activity for Scene 1 during the choice task when subjects rejected bad objects by *return* and *stay* and the neural activity when the objects were presented during the fixation task using GLMM. A GLMM was conducted to test the differences in neuronal activity for the target or saccade onset. The full and null models for the comparison are as follows:

*Full Model: NeuronalActivity* ∼ *Scene* × *Value* × *Direction* + (1|*monkey_ID*)

+ (1|*monkey_ID*: *Neuron_ID*),

*Null Model: NeuronalActivity* ∼ (1|*monkey_ID*) + (1|*monkey_ID*: *Neuron_ID*),

where *NeuronalActivity* is the mean PSTH of individual neurons during a 200 ms period, beginning 100 ms after target onset; *Scene* (1–4), *Value* (good vs. bad), and *Direction* (contralateral vs. ipsilateral) are fixed effects; and *monkey_ID* and *Neuron_ID* are random effects.

The statistical significance for the model comparison was set at α = 0.05. Considering that six pairwise t-tests were performed to compare the normalized neuronal activities for the good and bad objects during the choice task, statistical significance for the post-hoc test was set at α = 0.05/6 with Bonferroni correction.

## Results

### Behavior results in the choice task

During the choice task, the two monkeys demonstrated their ability to differentiate good from bad objects, as shown in Figures 2A and B. Raincloud plots in Figure 2A show the reaction times of both monkeys for good (saccade for accept) or bad objects (return go saccade for reject) in all four scenes. The cloud portion illustrates the distribution, and the raindrops represent individual data points. The median values and confidence intervals are highlighted in the boxplots (Allen et al., 2021). Table 1 summarizes the number of samples, mean, standard deviation (SD), and 95% confidence intervals (CI) of the reaction times toward good or bad objects in each *scene* in the neuronal recording sessions.

**Table 1.** Saccade reaction times in each condition of each monkey.

| **Monkey C** | **n** | **Mean RT (ms)** | **SD** | **95% CI** |
| --- | --- | --- | --- | --- |
| **Good obj** |  |  |  |  |
| ObjA in Scene 1 | 1428 | 160.9 | 21.6 | [159.7 to 162.0] |
| ObjC in Scene 2 | 1443 | 162.6 | 22.4 | [161.5 to 163.8] |
| ObjE in Scene 3 | 1452 | 169.4 | 22.1 | [168.3 to 170.5] |
| ObjF in Scene 4 | 1419 | 165.5 | 21.4 | [164.4 to 166.7] |
| **Bad obj** |  |  |  |  |
| ObjB in Scene 1 | 1076 | 234.3 | 46.7 | [231.5 to 237.1] |
| ObjD in Scene 2 | 1076 | 240.0 | 46.0 | [237.2 to 242.7] |
| ObjF in Scene 3 | 1247 | 215.0 | 50.3 | [212.2 to 217.8] |
| ObjE in Scene 4 | 1205 | 221.3 | 52.3 | [218.8 to 225.1] |
| **Monkey S** | **n** | **Mean RT (ms)** | **SD** | **95% CI** |
| **Good obj** |  |  |  |  |
| ObjA in Scene 1 | 1466 | 174.6 | 20.8 | [173.5 to 175.6] |
| ObjC in Scene 2 | 1482 | 179.1 | 22.0 | [178.0 to 180.2] |
| ObjE in Scene 3 | 1490 | 184.5 | 21.4 | [183.4 to 185.6] |
| ObjF in Scene 4 | 1476 | 186.9 | 24.3 | [185.7 to 188.2] |
| **Bad obj** |  |  |  |  |
| ObjB in Scene 1 | 1194 | 302.1 | 42.5 | [299.7 to 304.6] |
| ObjD in Scene 2 | 1054 | 311.9 | 41.2 | [309.4 to 314.4] |
| ObjF in Scene 3 | 1174 | 291.2 | 51.8 | [288.2 to 294.2] |
| ObjE in Scene 4 | 1120 | 299.9 | 47.4 | [297.1 to 302.6] |

Figure 2B quantifies the choices made by the monkeys when presented with bad objects. Table 2 lists the number of actions directed toward the bad objects in each *scene*. Monkeys consistently accepted good objects and frequently made a *return* (monkey C: 75.3%, 78.1%, 91.8%, and 85.1% in *scenes* 1, 2, 3, and 4, respectively; monkey S: 74.1%, 70.7%, 79.7%, and 78.2% in *scenes* 1, 2, 3, and 4, respectively) and *stay* (monkey C: 21.3%, 20.2%, 6.7%, and 11.3% in *scenes* 1, 2, 3, and 4, respectively; monkey S: 23.5%, 28.3%, 19.5%, and 20.1% in *scenes* 1, 2, 3, and 4, respectively) for bad objects. The target range was defined as 8° per side square during the task. When the monkeys made a saccade toward the target, their eye position moved out the range for < 400 ms, the target disappeared, and the fixation point appeared again.

**Table 2.** Counts of chosen actions for bad objects.

| **Monkey C** | Total | Accept | Return | Stay | Others | Fxbreak |
| --- | --- | --- | --- | --- | --- | --- |
| Scene 1 | 1479 | 0 | 1113 | 314 | 28 | 24 |
| Scene 2 | 1409 | 1 | 1100 | 284 | 15 | 9 |
| Scene 3 | 1396 | 6 | 1282 | 94 | 9 | 5 |
| Scene 4 | 1473 | 6 | 1253 | 167 | 13 | 34 |
| Non-switch (Scenes 1, 2) | 2888 | 1 | 2213 | 598 | 43 | 33 |
| Switch (Scenes 3, 4) | 2869 | 12 | 2535 | 261 | 22 | 39 |
| **Monkey S** | Total | Accept | Return | Stay | Others | Fxbreak |
| Scene 1 | 1602 | 3 | 1188 | 376 | 30 | 5 |
| Scene 2 | 1504 | 2 | 1063 | 426 | 6 | 7 |
| Scene 3 | 1475 | 1 | 1176 | 287 | 7 | 4 |
| Scene 4 | 1446 | 5 | 1130 | 291 | 15 | 5 |
| Non-switch (Scenes 1, 2) | 3106 | 5 | 2351 | 802 | 36 | 12 |
| Switch (Scenes 3, 4) | 2921 | 6 | 2306 | 578 | 22 | 9 |

Particularly, the monkeys had to wait for 400 ms for the next target to be presented if they rejected bad objects by *stay*, whereas the waiting time for the next target was shorter than 400 ms if they rejected bad objects by *return*. This may explain the frequent rejection of bad objects by *return*. The proportions of *stay* among the four scenes were significantly higher for *scenes* 1 and 2 than for *scenes* 3 and 4 for both monkeys (Fisher’s test, monkey C; *p* < 5.55×10^-36^, φ = 0.17, 95% CI: 0.32–0.45; monkey S: *p* < 1.49×10^-8^, φ = 0.07, 95% CI: 0.62–0.80). This may be attributed to the consistent object values for *scenes* 1 and 2 and the alternating values of the objects for *scenes* 3 and 4.

### Neuronal activity of striatal neurons during the choice task

To investigate the involvement of striatal neurons in object evaluation, neuronal activity was recorded from the caudate nucleus and putamen of monkeys undertaking the choice task. Through k-means clustering based on the responses to contralateral good and bad objects (Figure 3A), we identified three distinct clusters. Table 3 details the number of neurons in clusters 1 (n = 38), 2 (n = 47), and 3 (n = 53). Several task-related neurons were found in the anterior part of the striatum near the anterior commissure for both the caudate nucleus and putamen (Figure 3B).

**Table 3.**
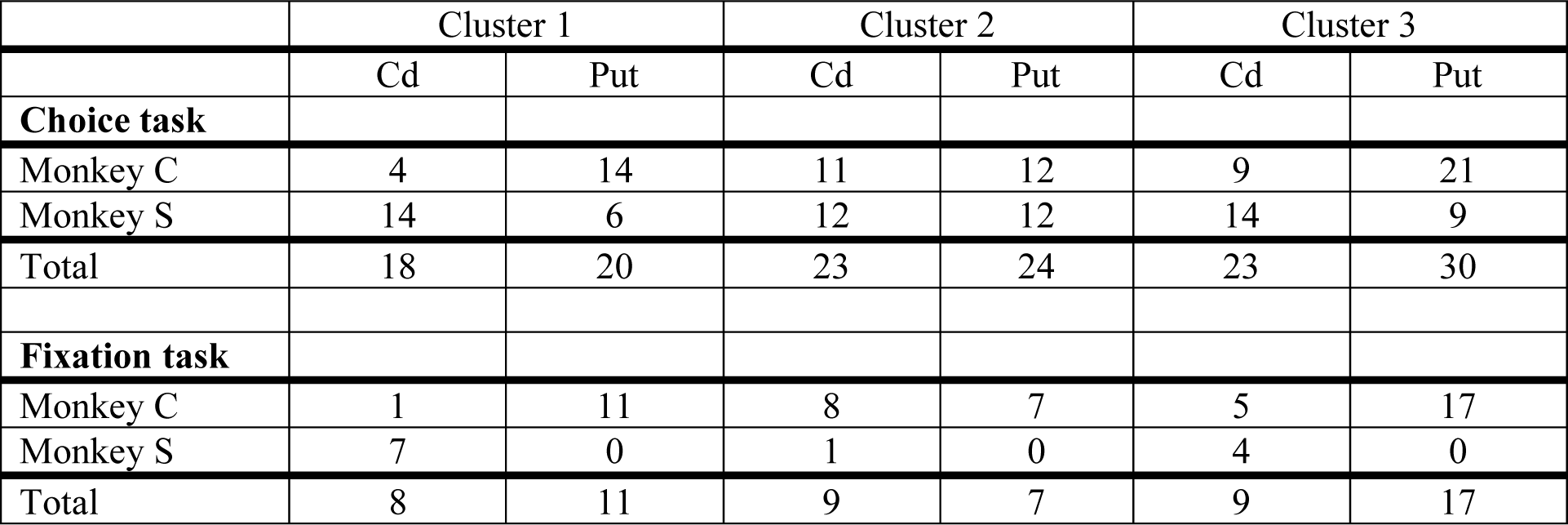
Numbers of recorded task-related neurons in the striatum of each monkey.

**Table 4.**
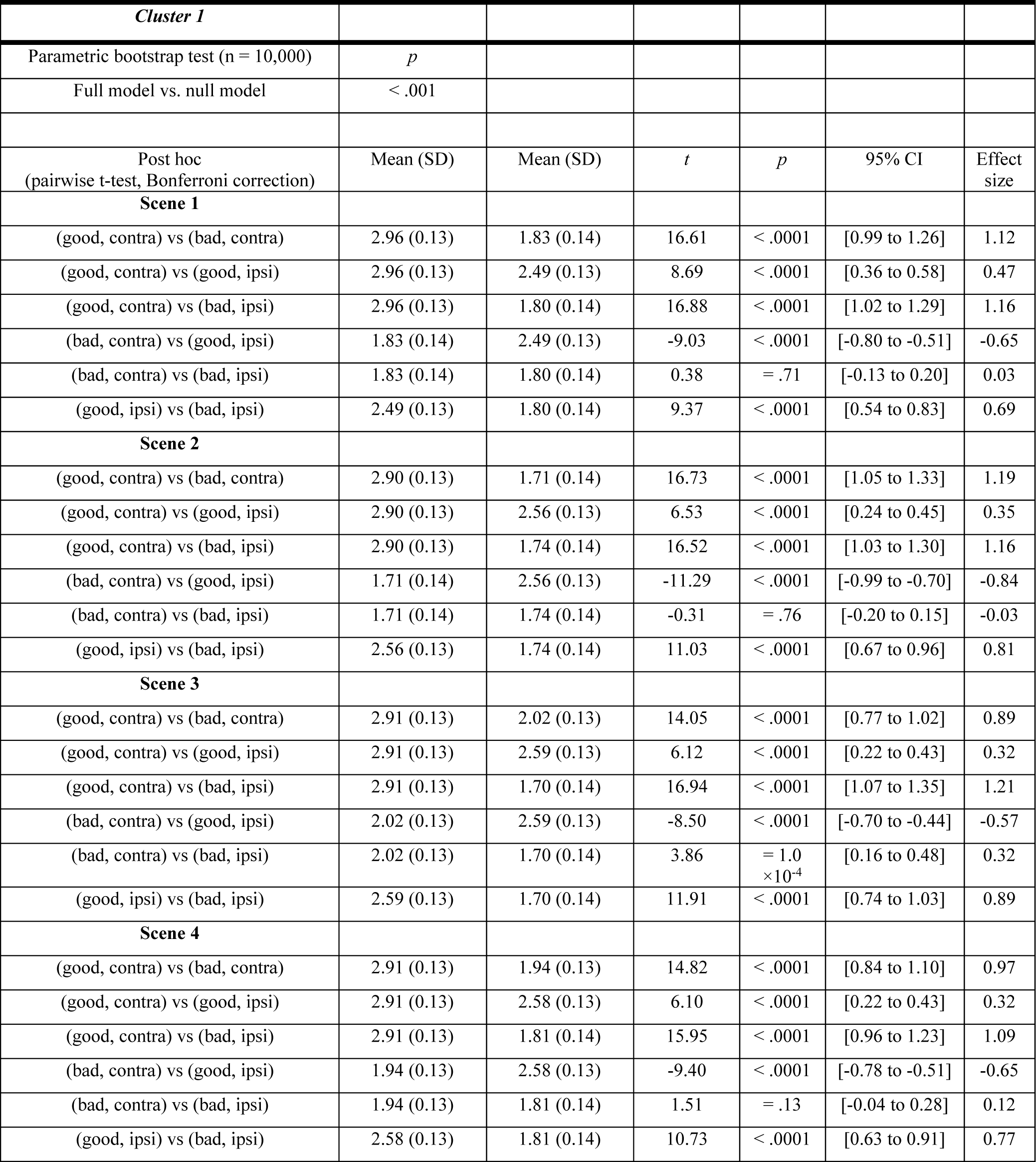
Summary of statistical test to compare the normalized neuronal activity of striatum neurons of cluster 1 at target onset among conditions during choice task in Figure 5.

| *Cluster 1* |  |  |  |  |  |  |
| --- | --- | --- | --- | --- | --- | --- |
| Parametric bootstrap test (n = 10,000) | *p* |  |  |  |  |  |
| Full model vs. null model | < .001 |  |  |  |  |  |
| Post hoc<br>(pairwise t-test, Bonferroni correction) | Mean (SD) | Mean (SD) | *t* | *p* | 95% CI | Effect size |
| **Scene 1** |  |  |  |  |  |  |
| (good, contra) vs (bad, contra) | 2.96 (0.13) | 1.83 (0.14) | 16.61 | < .0001 | [0.99 to 1.26] | 1.12 |
| (good, contra) vs (good, ipsi) | 2.96 (0.13) | 2.49 (0.13) | 8.69 | < .0001 | [0.36 to 0.58] | 0.47 |
| (good, contra) vs (bad, ipsi) | 2.96 (0.13) | 1.80 (0.14) | 16.88 | < .0001 | [1.02 to 1.29] | 1.16 |
| (bad, contra) vs (good, ipsi) | 1.83 (0.14) | 2.49 (0.13) | -9.03 | < .0001 | [-0.80 to -0.51] | -0.65 |
| (bad, contra) vs (bad, ipsi) | 1.83 (0.14) | 1.80 (0.14) | 0.38 | = .71 | [-0.13 to 0.20] | 0.03 |
| (good, ipsi) vs (bad, ipsi) | 2.49 (0.13) | 1.80 (0.14) | 9.37 | < .0001 | [0.54 to 0.83] | 0.69 |
| **Scene 2** |  |  |  |  |  |  |
| (good, contra) vs (bad, contra) | 2.90 (0.13) | 1.71 (0.14) | 16.73 | < .0001 | [1.05 to 1.33] | 1.19 |
| (good, contra) vs (good, ipsi) | 2.90 (0.13) | 2.56 (0.13) | 6.53 | < .0001 | [0.24 to 0.45] | 0.35 |
| (good, contra) vs (bad, ipsi) | 2.90 (0.13) | 1.74 (0.14) | 16.52 | < .0001 | [1.03 to 1.30] | 1.16 |
| (bad, contra) vs (good, ipsi) | 1.71 (0.14) | 2.56 (0.13) | -11.29 | < .0001 | [-0.99 to -0.70] | -0.84 |
| (bad, contra) vs (bad, ipsi) | 1.71 (0.14) | 1.74 (0.14) | -0.31 | = .76 | [-0.20 to 0.15] | -0.03 |
| (good, ipsi) vs (bad, ipsi) | 2.56 (0.13) | 1.74 (0.14) | 11.03 | < .0001 | [0.67 to 0.96] | 0.81 |
| **Scene 3** |  |  |  |  |  |  |
| (good, contra) vs (bad, contra) | 2.91 (0.13) | 2.02 (0.13) | 14.05 | < .0001 | [0.77 to 1.02] | 0.89 |
| (good, contra) vs (good, ipsi) | 2.91 (0.13) | 2.59 (0.13) | 6.12 | < .0001 | [0.22 to 0.43] | 0.32 |
| (good, contra) vs (bad, ipsi) | 2.91 (0.13) | 1.70 (0.14) | 16.94 | < .0001 | [1.07 to 1.35] | 1.21 |
| (bad, contra) vs (good, ipsi) | 2.02 (0.13) | 2.59 (0.13) | -8.50 | < .0001 | [-0.70 to -0.44] | -0.57 |
| (bad, contra) vs (bad, ipsi) | 2.02 (0.13) | 1.70 (0.14) | 3.86 | = 1.0<br>×10<sup>-4</sup> | [0.16 to 0.48] | 0.32 |
| (good, ipsi) vs (bad, ipsi) | 2.59 (0.13) | 1.70 (0.14) | 11.91 | < .0001 | [0.74 to 1.03] | 0.89 |
| **Scene 4** |  |  |  |  |  |  |
| (good, contra) vs (bad, contra) | 2.91 (0.13) | 1.94 (0.13) | 14.82 | < .0001 | [0.84 to 1.10] | 0.97 |
| (good, contra) vs (good, ipsi) | 2.91 (0.13) | 2.58 (0.13) | 6.10 | < .0001 | [0.22 to 0.43] | 0.32 |
| (good, contra) vs (bad, ipsi) | 2.91 (0.13) | 1.81 (0.14) | 15.95 | < .0001 | [0.96 to 1.23] | 1.09 |
| (bad, contra) vs (good, ipsi) | 1.94 (0.13) | 2.58 (0.13) | -9.40 | < .0001 | [-0.78 to -0.51] | -0.65 |
| (bad, contra) vs (bad, ipsi) | 1.94 (0.13) | 1.81 (0.14) | 1.51 | = .13 | [-0.04 to 0.28] | 0.12 |
| (good, ipsi) vs (bad, ipsi) | 2.58 (0.13) | 1.81 (0.14) | 10.73 | < .0001 | [0.63 to 0.91] | 0.77 |

**Table 5.** Summary of statistical test to compare the normalized neuronal activity of striatum neurons of cluster 2 at target onset among conditions during choice task in Figure 5.

| *Cluster 2* |  |  |  |  |  |  |
| --- | --- | --- | --- | --- | --- | --- |
| Parametric bootstrap test (n = 10,000) | *p* |  |  |  |  |  |
| Full model vs. null model | < .001 |  |  |  |  |  |
| Post hoc<br>(pairwise t-test, Bonferroni correction) | Mean (SD) | Mean (SD) | *t* | *p* | 95% CI | Effect size |
| **Scene 1** |  |  |  |  |  |  |
| (good, contra) vs (bad, contra) | 1.58 (0.14) | 2.43 (0.13) | -12.82 | < .0001 | [-0.98 to -0.72] | -0.71 |
| (good, contra) vs (good, ipsi) | 1.58 (0.14) | 1.63 (0.14) | -0.66 | = .51 | [-0.20 to 0.10] | 0.10 |
| (good, contra) vs (bad, ipsi) | 1.58 (0.14) | 2.50 (0.13) | -13.93 | < .0001 | [-1.04 to -0.78] | -0.77 |
| (bad, contra) vs (good, ipsi) | 2.43 (0.13) | 1.63 (0.14) | 12.27 | < .0001 | [0.67 to 0.92] | 0.93 |
| (bad, contra) vs (bad, ipsi) | 2.43 (0.13) | 1.80 (0.14) | -1.30 | = .19 | [-0.16 to 0.03] | 0.03 |
| (good, ipsi) vs (bad, ipsi) | 1.63 (0.14) | 2.50 (0.13) | -13.40 | < .0001 | [-0.99 to -0.73] | -0.73 |
| **Scene 2** |  |  |  |  |  |  |
| (good, contra) vs (bad, contra) | 1.75 (0.14) | 2.58 (0.13) | -13.59 | < .0001 | [-0.96 to -0.72] | -0.71 |
| (good, contra) vs (good, ipsi) | 1.75 (0.14) | 1.70 (0.14) | 0.66 | = .51 | [-0.10 to 0.19] | 0.19 |
| (good, contra) vs (bad, ipsi) | 1.75 (0.14) | 2.57 (0.13) | -13.35 | < .0001 | [-0.95 to -0.70] | -0.70 |
| (bad, contra) vs (good, ipsi) | 2.58 (0.13) | 1.70 (0.14) | 14.13 | < .0001 | [0.76 to 1.00] | 1.01 |
| (bad, contra) vs (bad, ipsi) | 2.58 (0.13) | 2.57 (0.13) | 0.20 | = .84 | [-0.09 to 0.11] | 0.11 |
| (good, ipsi) vs (bad, ipsi) | 1.70 (0.14) | 2.57 (0.13) | -13.90 | < .0001 | [-1.00 to -0.75] | -0.74 |
| **Scene 3** |  |  |  |  |  |  |
| (good, contra) vs (bad, contra) | 1.85 (0.14) | 2.54 (0.13) | -11.65 | < .0001 | [-0.81 to -0.57] | -0.57 |
| (good, contra) vs (good, ipsi) | 1.85 (0.14) | 1.75 (0.14) | 1.48 | = .14 | [-0.03 to 0.24] | 0.24 |
| (good, contra) vs (bad, ipsi) | 1.85 (0.14) | 2.45 (0.13) | -9.91 | < .0001 | [-0.72 to -0.48] | -0.48 |
| (bad, contra) vs (good, ipsi) | 2.54 (0.13) | 1.75 (0.14) | 12.94 | < .0001 | [0.67 to 0.91] | 0.92 |
| (bad, contra) vs (bad, ipsi) | 2.54 (0.13) | 2.45 (0.13) | 1.89 | = .06 | [0.00 to 0.19] | 0.19 |
| (good, ipsi) vs (bad, ipsi) | 1.75 (0.14) | 2.45 (0.13) | -11.24 | < .0001 | [-0.82 to -0.58] | -0.57 |
| **Scene 4** |  |  |  |  |  |  |
| (good, contra) vs (bad, contra) | 1.94 (0.14) | 2.65 (0.13) | -12.55 | < .0001 | [-0.82 to -0.60] | -0.59 |
| (good, contra) vs (good, ipsi) | 1.94 (0.14) | 1.93 (0.14) | 0.20 | = .84 | [-0.12 to 0.14] | 0.14 |
| (good, contra) vs (bad, ipsi) | 1.94 (0.14) | 2.64 (0.13) | -12.27 | < .0001 | [-0.81 to -0.58] | -0.58 |
| (bad, contra) vs (good, ipsi) | 2.65 (0.13) | 1.93 (0.14) | 12.72 | < .0001 | [0.61 to 0.83] | 0.84 |
| (bad, contra) vs (bad, ipsi) | 2.65 (0.13) | 2.64 (0.13) | 0.30 | = .76 | [-0.08 to 0.10] | 0.10 |
| (good, ipsi) vs (bad, ipsi) | 1.93 (0.14) | 2.64 (0.13) | -12.45 | < .0001 | [-0.82 to -0.60] | -0.59 |

Figure 4A shows a representative neuron from cluster 1. This neuron increased the firing rate more frequently when a good object was presented for all the scenes than when a bad object was. In contrast, one representative neuron in cluster 2 exclusively increased the firing rate in response to bad objects (Figure 4B). The example neuron of cluster 3 showed a similar response only after the target onset for the good and bad objects, and the neuronal activity for the bad objects was higher than that for the good objects.

**Figure 4.**
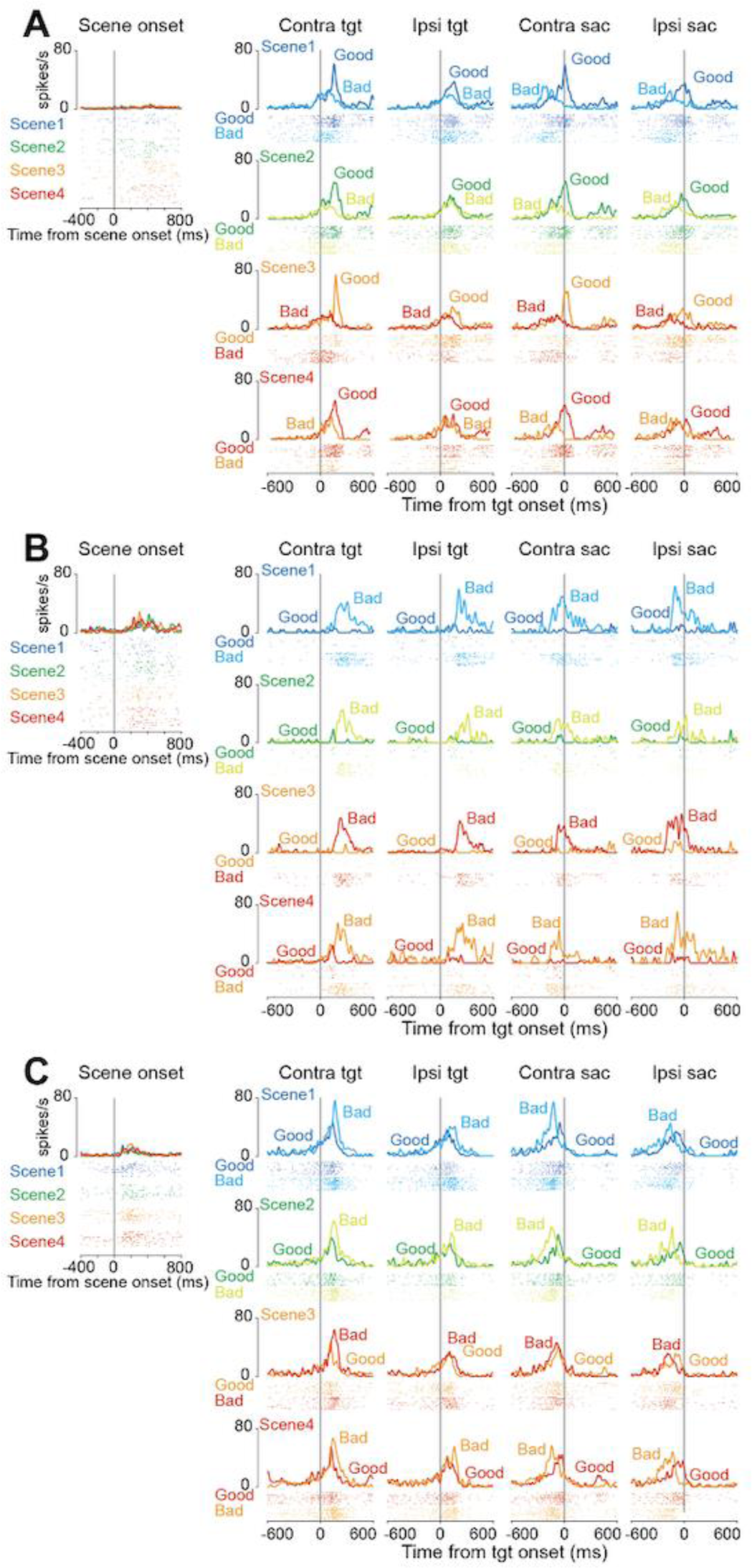
Representative neurons of the three clusters. **(A, B, C)** The raster plots and histograms of representative neurons of clusters 1 **(A)**, 2 **(B)**, and 3 **(C)**. Data are aligned with the scene onset, target onset, or saccade initiation (vertical line in each panel). Traces in different colors indicate the spike density for good or bad objects in each scene.

Figure 5 shows the population activity of the three clusters at the scene (A, E, and I) and target onset (C, F, and J). The lower panels illustrate the normalized neuronal activities of individual neurons in the three clusters at scene onset (scenes 1-4) and on the presentations of good and bad objects. The violin plots in Figures 5B, D, G, H, K, and I show the mean firing rate of individual neurons during the 200 ms from 100 to 300 ms after the scene or target onset. For quantitative analysis, we conducted a parametric bootstrap test for the GLMM and subsequent post-hoc pairwise t-tests with Bonferroni correction (see the Methods section for details).

**Figure 5.**
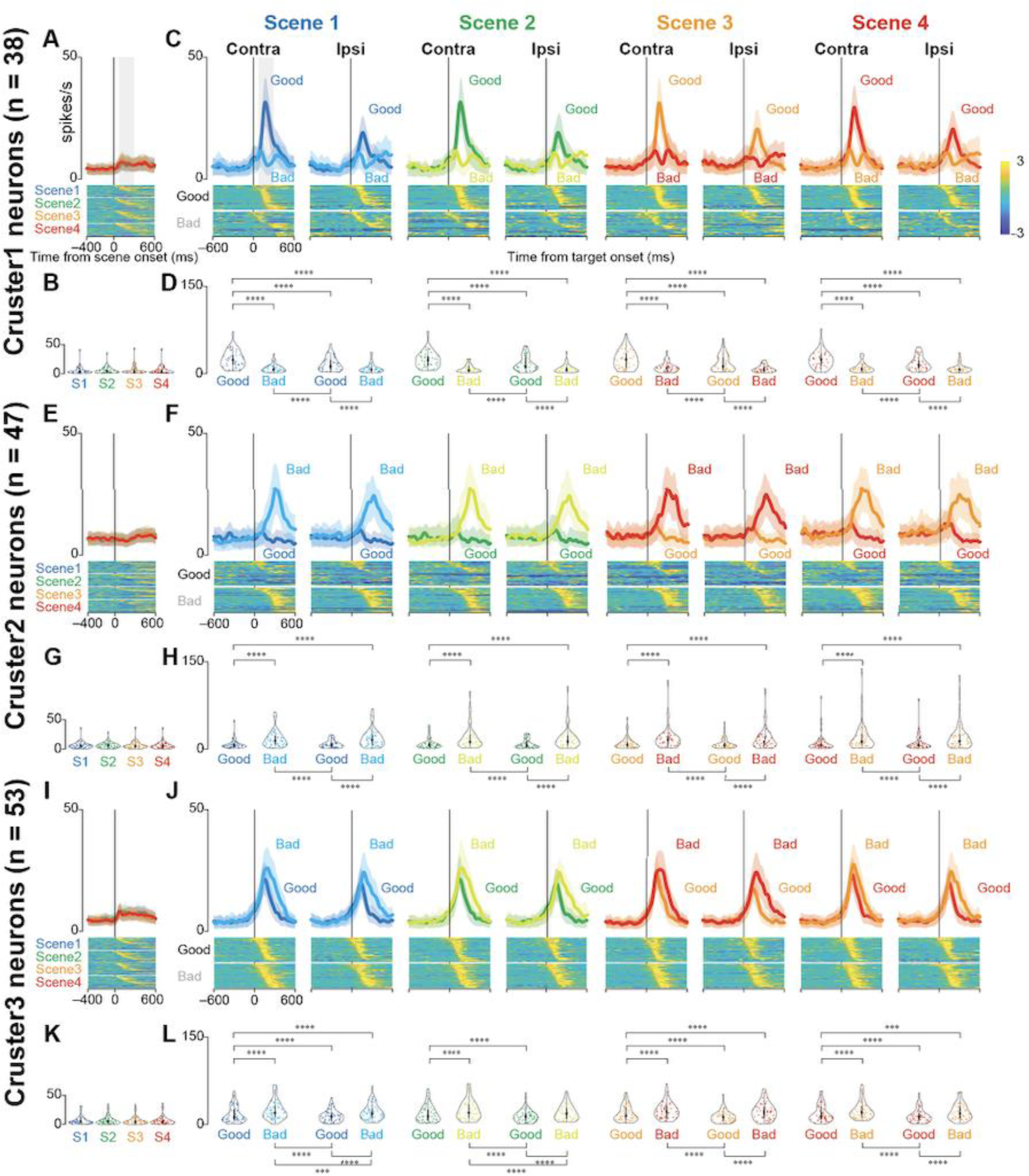
Population activity of three groups of striatal neurons at scene and target onsets during the choice task. **(A, E, and I)** Mean population activities aligned at scene onset (scenes 1–4) during the choice task for the neurons in clusters 1 **(A)**, 2 **(E)**, and 3 **(I)**. The shaded areas indicate ± SEMs. The lower panels show the color maps of the normalized firing rates of individual neurons. **(B, G, and K)** The violin plots of the mean firing rates of individual neurons in clusters 1 **(B)**, 2 **(G)**, and 3 **(K)** for each scene onset during the choice task. Neuronal activity is measured for a 200-ms interval beginning from 100 ms after scene onset (gray rectangle in **[A]**). The larger circle indicates the median value, the thick vertical line shows the range between the first and the third quartile, and the thin vertical line indicates the range from the lower to the upper adjacent values. **(C, F, and J)** Mean population activities aligned at the contralateral and ipsilateral good and bad target onsets for scenes 1–4 during the choice task of the neurons in clusters 1 **(C)**, 2 **(F)**, and 3 **(J)**. The shaded areas indicate ± SEMs (standard error of the mean). The lower panels show the color maps of the normalized firing rates of individual neurons. **(D, H, and L)** The violin plots of the mean firing rates of individual neurons in clusters 1 **(B)**, 2 **(G)**, and 3 **(K)** for each scene onset during the choice task. Neuronal activity is measured for a 200-ms interval beginning from 100 ms after target onset (gray rectangle in the panel of contralateral target onset in Scene 1 in **[C]**). The asterisk indicates a significant difference in neuronal activity among the conditions in the choice and fixation tasks (post-hoc pairwise t-tests with Bonferroni correction, \**p <* 0.05, \*\**p* < 0.01, \*\*\**p* < 0.001, \*\*\*\**p* < 0.0001). Asterisks are only attached to combinations with significant differences.

Statistical analysis showed significant differences in neuronal activity when objects were presented during the choice task (post-hoc pairwise t-tests, *p* < 0.0001). The detailed results of the other post-hoc pairwise test comparisons are summarized in Tables 4 (cluster 1), 5 (cluster 2), and 6 (cluster 3). There were no significant differences in neuronal activity at the onsets of scenes 1–4 (parametric bootstrap tests, full model vs. null models; *p* = 0.69 (cluster 1), *p* = 0.98 (cluster 2), and *p* = 0.35 (cluster 3).

Neurons in cluster 1 were more active when presented with a good object than when presented with a bad object. They were more active when presented with a good object on their contralateral side, suggesting that they were involved in accepting the good object. In contrast, neurons in cluster 2 were more active when presented with a bad object, suggesting that they were involved in rejecting the bad object. The cluster 3 neurons showed changes in neural activity immediately after the target was presented, and the temporal changes in neuronal activity immediately after the target was presented were similar for good and bad objects. This suggested that they were involved in the visual response. The changes in neural activity in response to bad objects lasted longer. This may be attributed to the faster saccade reaction times; therefore, the visual response to the good objects was attenuated more quickly.

To examine whether the neuronal activity observed during the choice task reflected the visual characteristics or values of the objects presented, we used a set of four scene conditions (*scenes* 1–4). There were no significant differences in neuronal activity across scenes. These results suggest that the differences in neural activity observed during target onset are not due to differences in the visual characteristics of the target, but rather differences in the values of the reward for the target.

### Negative correlation between neuronal activity and saccade reaction times

Statistical tests were also performed with data aligned at saccade initiation (Figure 6). There was no obvious difference in the population activity for all three clusters at target onset across the four scene conditions. Therefore, the merged data from Scenes 1, 2, 3, and 4 were used for this analysis. Figures 6A, D, and G show the population activity aligned at the saccade for choosing good targets (accept) or saccades for rejecting bad targets (return go). GLMM and subsequent post hoc pairwise t-tests revealed significant differences in neuronal activity when the objects were presented during the choice task. Detailed results of other post hoc pairwise test comparisons are summarized in Table 7.

**Figure 6.**
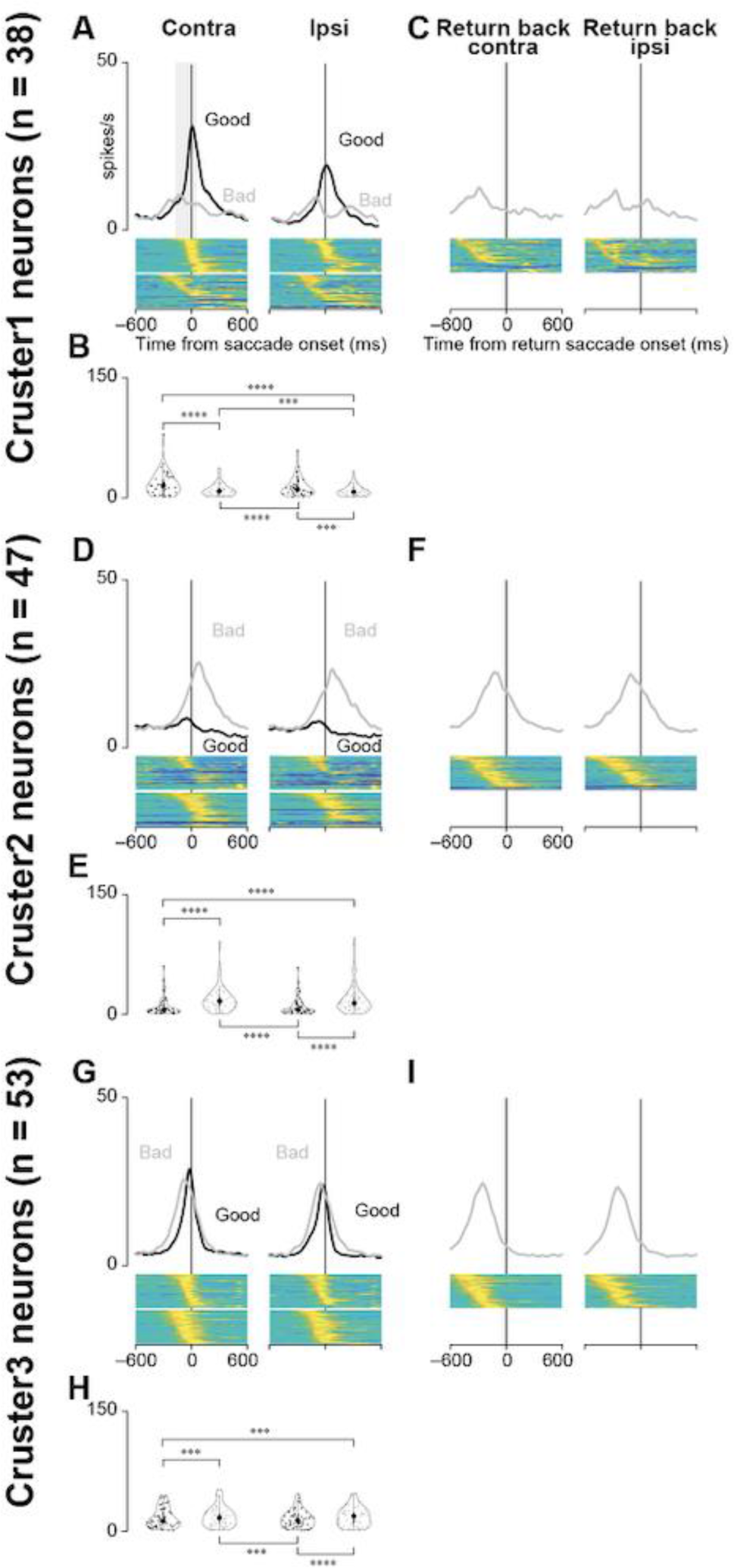
Population activity of three groups of striatal neurons at saccade initiation during the choice task. **(A, D, and G)** Mean population activities aligned at saccade initiation toward contralateral or ipsilateral good or bad objects in all scenes during the choice task for the neurons in clusters 1 **(A)**, 2 **(D)**, and 3 **(G)**. The shaded areas indicate ± SEMs. The lower panels show the color maps of the normalized firing rates of individual neurons. **(B, E, and H)** The violin plots of the mean firing rates of individual neurons in clusters 1 **(B)**, 2 **(E)**, and 3 **(G)** when the monkeys make a saccade to the target during the choice task. Neuronal activity is measured for a 200-ms interval from 150 ms before the saccade initiation (gray rectangle in **(A)**). The asterisk indicates a significant difference in neuronal activity among the conditions for the choice tasks (post-hoc pairwise t-tests with Bonferroni correction, \**p <* 0.05, \*\**p* < 0.01, \*\*\**p* < 0.001, \*\*\*\**p* < 0.0001). Asterisks are attached only to combinations with significant differences. **(C, F, and I)** Mean population activities aligned at the initiation of the return saccade after the monkeys make a saccade to bad objects (clusters 1 **[C]**, 2 **[F]**, and 3 **[I]**).

**Table 6.** Summary of statistical test to compare the normalized neuronal activity of striatum neurons of cluster3 at target onset among conditions during choice task in Figure 5.

| *Cluster 3* |  |  |  |  |  |  |
| --- | --- | --- | --- | --- | --- | --- |
| Parametric bootstrap test (n = 10,000) | *p* |  |  |  |  |  |
| Full model vs. null model | < .001 |  |  |  |  |  |
| Post hoc<br>(pairwise t-test, Bonferroni correction) | Mean (SD) | Mean (SD) | *t* | *p* | 95% CI | Effect size |
| **Scene 1** |  |  |  |  |  |  |
| (good, contra) vs (bad, contra) | 2.66 (0.09) | 3.00 (0.08) | -7.86 | < .0001 | [-0.43 to -0.26] | -0.35 |
| (good, contra) vs (good, ipsi) | 2.66 (0.09) | 2.42 (0.07) | 4.69 | < .0001 | [0.14 to 0.34] | 0.24 |
| (good, contra) vs (bad, ipsi) | 2.66 (0.09) | 2.86 (0.08) | -4.50 | < .0001 | [-0.29 to -0.12] | -0.20 |
| (bad, contra) vs (good, ipsi) | 3.00 (0.08) | 2.42 (0.07) | 12.33 | < .0001 | [0.49 to 0.68] | 0.59 |
| (bad, contra) vs (bad, ipsi) | 3.00 (0.08) | 2.86 (0.08) | 3.41 | = 7.0<br>×10<sup>-4</sup> | [0.06 to 0.22] | 0.14 |
| (good, ipsi) vs (bad, ipsi) | 2.42 (0.07) | 2.86 (0.08) | -9.10 | < .0001 | [-0.54 to -0.35] | -0.44 |
| **Scene 2** |  |  |  |  |  |  |
| (good, contra) vs (bad, contra) | 2.68 (0.08) | 3.00 (0.08) | -7.32 | < .0001 | [-0.41 to -0.23] | -0.32 |
| (good, contra) vs (good, ipsi) | 2.68 (0.08) | 2.44 (0.09) | 4.88 | < .0001 | [0.15 to 0.34] | 0.25 |
| (good, contra) vs (bad, ipsi) | 2.68 (0.08) | 2.81 (0.08) | -2.67 | = 7.6<br>×10<sup>-3</sup> | [-0.21 to -0.03] | -0.12 |
| (bad, contra) vs (good, ipsi) | 3.00 (0.08) | 2.44 (0.09) | 12.00 | < .0001 | [0.47 to 0.66] | 0.56 |
| (bad, contra) vs (bad, ipsi) | 3.00 (0.08) | 2.81 (0.08) | 4.69 | < .0001 | [0.12 to 0.28] | 0.20 |
| (good, ipsi) vs (bad, ipsi) | 2.44 (0.09) | 2.81 (0.08) | -7.50 | < .0001 | [-0.46 to -0.27] | -0.37 |
| **Scene 3** |  |  |  |  |  |  |
| (good, contra) vs (bad, contra) | 2.72 (0.08) | 2.99 (0.08) | -6.14 | < .0001 | [-0.35 to -0.17] | -0.27 |
| (good, contra) vs (good, ipsi) | 2.72 (0.08) | 2.38 (0.09) | 6.82 | < .0001 | [0.25 to 0.45] | 0.35 |
| (good, contra) vs (bad, ipsi) | 2.72 (0.08) | 2.88 (0.08) | -4.10 | < .0001 | [-0.27 to -0.09] | -0.18 |
| (bad, contra) vs (good, ipsi) | 2.99 (0.08) | 2.38 (0.09) | 12.71 | < .0001 | [0.52 to 0.71] | 0.61 |
| (bad, contra) vs (bad, ipsi) | 2.99 (0.08) | 2.88 (0.08) | 2.07 | = 3.9<br>×10<sup>-2</sup> | [0.00 to 0.17] | 0.09 |
| (good, ipsi) vs (bad, ipsi) | 2.38 (0.09) | 2.88 (0.08) | -10.77 | < .0001 | [-0.62 to -0.43] | -0.53 |
| **Scene 4** |  |  |  |  |  |  |
| (good, contra) vs (bad, contra) | 2.73 (0.08) | 2.99 (0.08) | -5.95 | < .0001 | [-0.34 to -0.17] | -0.26 |
| (good, contra) vs (good, ipsi) | 2.73 (0.08) | 2.47 (0.09) | 5.29 | < .0001 | [0.16 to 0.36] | 0.26 |
| (good, contra) vs (bad, ipsi) | 2.73 (0.08) | 2.88 (0.08) | -3.28 | = 1.1<br>×10<sup>-3</sup> | [-0.23 to -0.06] | -0.15 |
| (bad, contra) vs (good, ipsi) | 2.99 (0.08) | 2.47 (0.09) | 11.08 | < .0001 | [0.43 to 0.61] | 0.52 |
| (bad, contra) vs (bad, ipsi) | 2.99 (0.08) | 2.88 (0.08) | 2.70 | = 6.9<br>×10<sup>-3</sup> | [0.03 to 0.19] | 0.11 |
| (good, ipsi) vs (bad, ipsi) | 2.47 (0.09) | 2.88 (0.08) | -8.49 | < .0001 | [-0.50 to -0.31] | -0.41 |

**Table 7.** Summary of statistical test to compare the mean neuronal activity of striatum neurons of clusters 1, 2, and 3 at saccade onset among conditions during choice task in Figure 6.

| *Cluster 1* |  |  |  |  |  |  |
| --- | --- | --- | --- | --- | --- | --- |
| Parametric bootstrap test (n = 10,000) | *p* |  |  |  |  |  |
| Full model vs. null model | < .001 |  |  |  |  |  |
| Post hoc<br>(pairwise t-test, Bonferroni correction) | Mean (SD) | Mean (SD) | *t* | *p* | 95% CI | Effect size |
| (good, contra) vs (bad, contra) | 2.38 (0.05) | 1.77 (0.07) | 7.41 | < .0001 | [0.45 to 0.78] | 0.61 |
| (good, contra) vs (good, ipsi) | 2.38 (0.05) | 2.26 (0.06) | 1.76 | = $7.9 \times 10^{-2}$ | [-0.01 to 0.27] | 0.13 |
| (good, contra) vs (bad, ipsi) | 2.38 (0.05) | 2.03 (0.06) | 4.52 | < .0001 | [0.20 to 0.50] | 0.35 |
| (bad, contra) vs (good, ipsi) | 1.77 (0.07) | 2.26 (0.06) | -5.76 | < .0001 | [-0.65 to -0.32] | -0.49 |
| (bad, contra) vs (bad, ipsi) | 1.77 (0.07) | 2.03 (0.06) | -2.99 | = $2.8 \times 10^{-3}$ | [-0.44 to -0.09] | -0.27 |
| (good, ipsi) vs (bad, ipsi) | 2.26 (0.06) | 2.03 (0.06) | 2.86 | = $4.5 \times 10^{-3}$ | [0.07 to 0.38] | 0.22 |
| *Cluster 2* |  |  |  |  |  |  |
| Parametric bootstrap test (n = 10,000) | *p* |  |  |  |  |  |
| Full model vs. null model | < .001 |  |  |  |  |  |
| Post hoc<br>(pairwise t-test, Bonferroni correction) | Mean (SD) | Mean (SD) | *t* | *p* | 95% CI | Effect size |
| (good, contra) vs (bad, contra) | 1.58 (0.07) | 2.85 (0.04) | -17.08 | < .0001 | [-1.41 to -1.12] | -1.27 |
| (good, contra) vs (good, ipsi) | 1.58 (0.07) | 1.47 (0.07) | 1.15 | = .25 | [-0.08 to 0.30] | 0.11 |
| (good, contra) vs (bad, ipsi) | 1.58 (0.07) | 2.85 (0.04) | -17.06 | < .0001 | [-1.41 to -1.12] | -1.27 |
| (bad, contra) vs (good, ipsi) | 2.85 (0.04) | 1.47 (0.07) | 17.77 | < .0001 | [1.22 to 1.53] | 1.38 |
| (bad, contra) vs (bad, ipsi) | 2.85 (0.04) | 2.85 (0.04) | 0.03 | = .98 | [-0.10 to 0.10] | 0.00 |
| (good, ipsi) vs (bad, ipsi) | 1.47 (0.07) | 2.85 (0.04) | -17.75 | < .0001 | [-1.53 to -1.22] | -1.38 |
| *Cluster 3* |  |  |  |  |  |  |
| Parametric bootstrap test (n = 10,000) | *p* |  |  |  |  |  |
| Full model vs. null model | < .001 |  |  |  |  |  |
| Post hoc<br>(pairwise t-test, Bonferroni correction) | Mean (SD) | Mean (SD) | *t* | *p* | 95% CI | Effect size |
| (good, contra) vs (bad, contra) | 1.57 (0.15) | 1.96 (0.15) | -2.95 | = $3.2 \times 10^{-3}$ | [-0.41 to -0.08] | -0.25 |
| (good, contra) vs (good, ipsi) | 1.57 (0.15) | 1.32 (0.16) | 2.68 | = $7.4 \times 10^{-3}$ | [-0.07 to 0.44] | 0.25 |
| (good, contra) vs (bad, ipsi) | 1.57 (0.15) | 1.71 (0.14) | -4.77 | < .0001 | [-5.44 to -0.28] | -0.39 |
| (bad, contra) vs (good, ipsi) | 1.96 (0.15) | 1.32 (0.16) | 5.55 | < .0001 | [0.32 to 0.67] | 0.50 |
| (bad, contra) vs (bad, ipsi) | 1.96 (0.15) | 1.71 (0.14) | -1.85 | = .06 | [-0.29 to 0.01] | -0.14 |
| (good, ipsi) vs (bad, ipsi) | 1.32 (0.16) | 1.71 (0.14) | -7.29 | < .0001 | [-0.81 to -0.47] | -0.64 |

Figures 6C, F, and I illustrate the population activity aligned at the initiation of return saccades (from the target to the center) when the monkeys rejected bad objects. The modulation of neural activity in cluster 1 neurons was minimal. The mean neural activities for cluster 2 and 3 neurons for bad objects started to decrease well before the initiation of the “return back” saccade. This suggests that the average neural activity of cluster 2 and 3 neurons is more related to the “return go” (initial saccade toward bad objects) or visual response than to the “return back” saccade.

This study examined the role of striatal neurons in saccade facilitation. Neuronal data were sorted into quantiles based on saccade latencies and stratified according to the value and direction of object presentation. Figures 7A, D, and G show the collective neural activity for these groups at the target presentation. The correlation between the mean normalized firing rate and saccade latencies for individual neurons is graphically represented in Figures 7B, E, and H Figures 7C, F, and I show histograms of the correlation coefficients for each neuron. Notably, the median correlation coefficient for cluster 1 neurons, when presented with contralateral good objects, was significantly less than zero (Wilcoxon signed-rank test, *p* = 0.02), suggesting that these neurons influenced the expedited initiation of saccades toward rewarding stimuli presented contralaterally.

**Figure 7.**
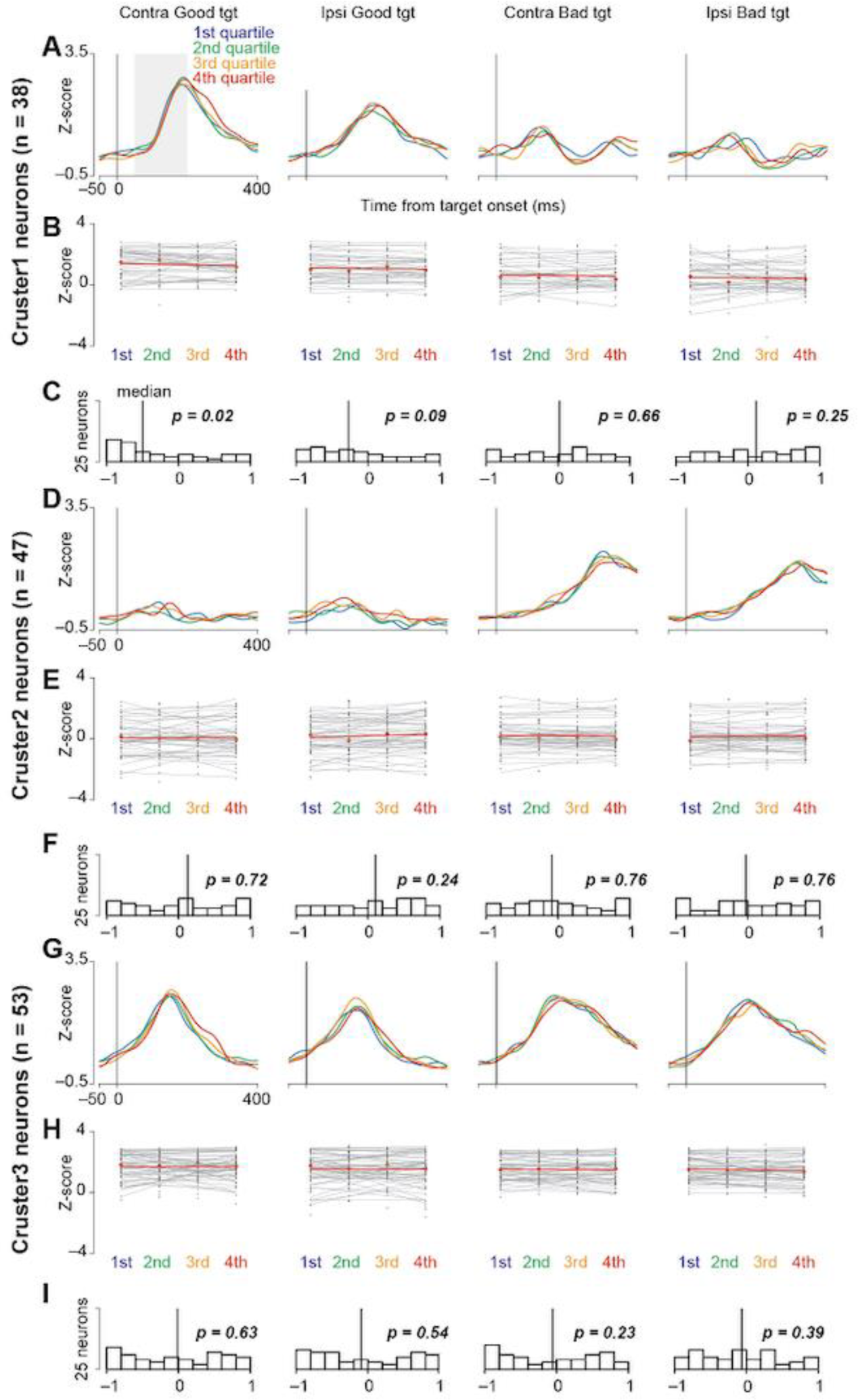
Correlation between neuronal activity and saccade reaction time. **(A, D, and G)** Time courses of the population activity of neurons in clusters 1 **(A)**, 2 **(D)**, and 3 **(G)** when the monkeys make a saccade toward bilateral good or bad targets. For individual neurons, the data are divided into four groups with an equal number of trials according to the saccade reaction times. The population activity of these four groups is illustrated in the differently colored lines. **(B, E, and H)** The regression slopes calculated from the mean neuronal activity classified into four groups according to saccade reaction times for each neuron in clusters 1 **(B)**, 2 **(E)**, and 3 **(G)**. Neuronal activity is measured for a 150-ms interval from 50 ms after the target onset (gray rectangle in **[A]**). **(C, F, and I)** Histograms of the distributions of coefficients computed for neuronal activity and individual saccade reaction times (clusters 1 **[C]**, 2 **[F]**, and 3 **[I]**). The vertical lines indicate the median values of each histogram.

### Activated striatal neurons while rejecting the choice not in simple suppressing saccades

Our analysis further probed activation patterns of the striatal neurons during active rejection of choices, distinguishing between “return” (making a saccade toward but not fixating on the bad object) and “stay” (maintaining gaze near the center point), as shown in Figure 8A. This differentiation was crucial as it indicated whether neuronal activity merely reflected saccade suppression or a more complex cognitive process. Utilizing the fixation task (Figure 8B) for neurons that exhibited modulated activity during the choice task, we implemented a parametric bootstrap test for a GLMM and conducted post hoc pairwise t-tests with Bonferroni correction to compare the activities across conditions.

**Figure 8.**
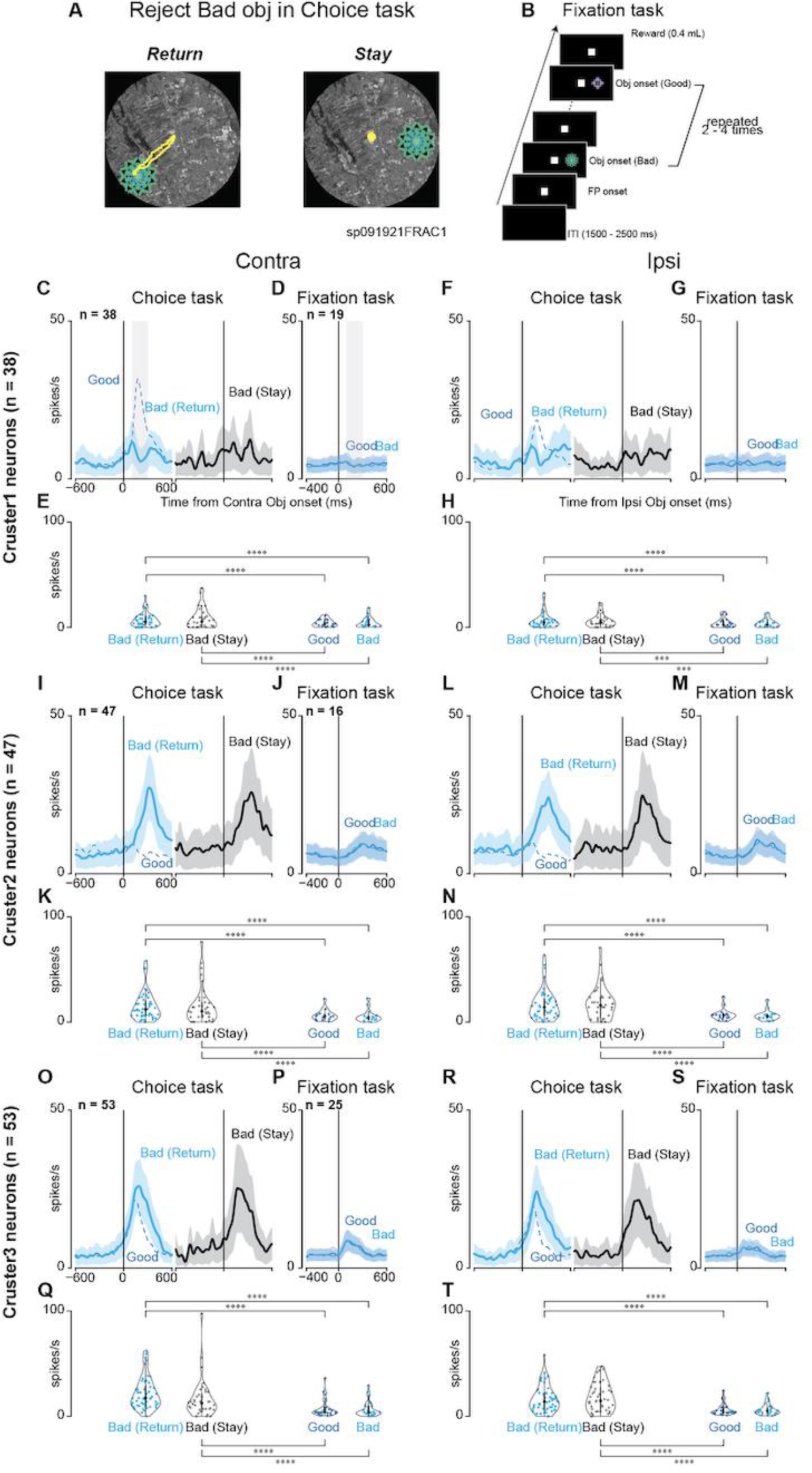
Comparison between return and stay when the monkeys rejected bad objects. **(A)** Examples of eye tracing when a monkey rejects a bad object by making a return saccade (Return) and fixating at the center point (Stay). **(B)** The procedure for the fixation task. During this task, good and bad objects with stable values used in scene 1 during the choice task are presented on the contralateral or ipsilateral side two to four times in a row. A reward is given at the end if a monkey could keep looking at the center point, suppressing reflexive saccade to the presented object. **(C, F, I, L, O, and R)** Mean population activities when the monkeys rejected bad objects in scene 1 by “return” (cyan-colored line) or “stay” (black-colored line) on the contralateral (clusters 1 **[C]**, 2 **[I]**, and 3 **[O]**) or ipsilateral (clusters 1 **[F]**, 2 **[L]**, and 3 **[R]**) side. For comparison, the blue dot line represents the mean population activity when the monkeys accept good objects in scene 1. **(D, G, J, M, P, and S)** Mean population activities when the monkeys suppressed the reflexive saccade to the presented good objects (blue-colored line) or bad objects (cyan-colored line) at the contralateral (clusters 1 **[D]**, 2 **[J]**, and 3 **[P]**) or ipsilateral (clusters 1 **[G]**, 2 **[M]**, and 3 **[S]**) side during the fixation task. **(E, H, K, N, Q, and T)** The violin plots of the mean firing rates of individual neurons during the choice task (clusters 1 **[E]**, 2 **[K]**, and 3 **[Q]**) and the fixation task (clusters 1 **[H]**, 2 **[N]**, and 3 **[T]**). Neuronal activity is measured for a 200-ms interval from 100 ms after object onset (gray rectangle in the panel of contralateral target onset in **[C]** or **[D]**). The asterisk indicates a significant difference in neuronal activity among the conditions for the tasks (post-hoc pairwise t-tests with Bonferroni correction, \**p <* 0.05, \*\**p* < 0.01, \*\*\**p* < 0.001, \*\*\*\**p* < 0.0001). Asterisks are attached only to combinations with significant differences.

No notable differences were found between the neuronal activities for the “return” and “stay” decisions or the good and bad objects during the fixation task. However, the observed modulation of neurons in clusters 1–3 during the choice task diminished during the fixation task. This attenuation highlights a potential difference in the roles of neurons. Although their activity decreased during the reactive inhibition required during the fixation task, it was more pronounced during the choice task when proactive inhibition was necessary. These findings, as summarized in Table 8, suggest that neurons in cluster 2 are particularly active during the inhibition process, as they show similar activity patterns irrespective of “return” or “stay” decisions. The reduced activation during the fixation task implied that these neurons were less involved in simple motor suppression and more involved in complex goal-directed behaviors. This distinction underscores the potential role of striatal neurons in proactive inhibition, wherein they contribute to the selection or rejection of actions to achieve the desired outcome.

**Table 8.** Summary of statistical test to compare the normalized neuronal activity of striatum neurons of clusters 1, 2, and 3 among Return, Stay during choice task, and fixation task in Figure 8.

| *Cluster1* |  |  |  |  |  |  |
| --- | --- | --- | --- | --- | --- | --- |
| Parametric bootstrap test (n = 10,000) | *p* |  |  |  |  |  |
| Full model vs. null model | < .001 |  |  |  |  |  |
| Post hoc<br>(pairwise t-test, Bonferroni correction) | Mean (SD) | Mean (SD) | *t* | *p* | 95% CI | Effect size |
| Contra |  |  |  |  |  |  |
| (Return, choice) vs (Stay, choice) | 1.73 (0.15) | 1.80 (0.15) | -0.90 | = .37 | [-0.24 to 0.09] | -0.08 |
| (Return, choice) vs (Good, fixation) | 1.73 (0.15) | 1.06 (0.17) | 5.46 | < .0001 | [0.42 to 0.90] | 0.66 |
| (Return, choice) vs (Bad, fixation) | 1.73 (0.15) | 1.11 (0.17) | 5.19 | < .0001 | [0.38 to 0.85] | 0.62 |
| (Stay, choice) vs (Good, fixation) | 1.80 (0.15) | 1.06 (0.17) | 5.99 | < .0001 | [0.50 to 0.98] | 0.74 |
| (Stay, choice) vs (Bad, fixation) | 1.80 (0.15) | 1.11 (0.17) | 5.73 | < .0001 | [0.46 to 0.93] | 0.69 |
| (Good, fixation) vs (Bad, fixation) | 1.06 (0.17) | 1.11 (0.17) | -0.29 | = .77 | [-0.33 to 0.24] | -0.04 |
| Ipsi |  |  |  |  |  |  |
| (Return, choice) vs (Stay, choice) | 1.69 (0.15) | 1.50 (0.15) | 2.14 | = .03 | [0.02 to 0.38] | 0.20 |
| (Return, choice) vs (Good, fixation) | 1.69 (0.15) | 0.98 (0.18) | 5.66 | < .0001 | [0.46 to 0.95] | 0.71 |
| (Return, choice) vs (Bad, fixation) | 1.69 (0.15) | 1.00 (0.17) | 5.52 | < .0001 | [0.44 to 0.93] | 0.68 |
| (Stay, choice) vs (Good, fixation) | 1.50 (0.15) | 0.98 (0.18) | 3.89 | < .001 | [0.25 to 0.77] | 0.51 |
| (Stay, choice) vs (Bad, fixation) | 1.50 (0.15) | 1.00 (0.17) | 3.74 | < .001 | [0.23 to 0.74] | 0.49 |
| (Good, fixation) vs (Bad, fixation) | 0.98 (0.18) | 1.00 (0.17) | -0.15 | = .87 | [-0.32 to 0.27] | -0.02 |
| *Cluster 2* |  |  |  |  |  |  |
| Parametric bootstrap test (n = 10,000) | *p* |  |  |  |  |  |
| Full model vs. null model | < .001 |  |  |  |  |  |
| Post hoc<br>(pairwise t-test, Bonferroni correction) | Mean (SD) | Mean (SD) | *t* | *p* | 95% CI | Effect size |
| Contra |  |  |  |  |  |  |
| (Return, choice) vs (Stay, choice) | 2.45 (0.14) | 2.49 (0.14) | -0.81 | = .42 | [-0.15 to 0.06] | -0.04 |
| (Return, choice) vs (Good, fixation) | 2.45 (0.14) | 1.50 (0.17) | 8.94 | < .0001 | [0.75 to 1.16] | 0.95 |
| (Return, choice) vs (Bad, fixation) | 2.45 (0.14) | 1.50 (0.17) | 8.94 | < .0001 | [0.75 to 1.16] | 0.95 |
| (Stay, choice) vs (Good, fixation) | 2.49 (0.14) | 1.50 (0.17) | 9.20 | < .0001 | [0.79 to 1.21] | 0.99 |
| (Stay, choice) vs (Bad, fixation) | 2.49 (0.14) | 1.50 (0.17) | 9.20 | < .0001 | [0.79 to 1.21] | 0.99 |
| (Good, fixation) vs (Bad, fixation) | 1.50 (0.17) | 1.50 (0.17) | 0.00 | = .99 | [-0.27 to 0.27] | 0.00 |
| Ipsi |  |  |  |  |  |  |
| (Return, choice) vs (Stay, choice) | 2.51 (0.14) | 2.52 (0.14) | -0.81 | = .82 | [-0.11 to 0.09] | -0.01 |
| (Return, choice) vs (Good, fixation) | 2.51 (0.14) | 1.59 (0.16) | 8.94 | < .0001 | [0.73 to 1.13] | 0.93 |
| (Return, choice) vs (Bad, fixation) | 2.51 (0.14) | 1.60 (0.16) | 8.94 | < .0001 | [0.71 to 1.11] | 0.91 |
| (Stay, choice) vs (Good, fixation) | 2.52 (0.14) | 1.59 (0.16) | 9.20 | < .0001 | [0.73 to 1.14] | 0.94 |
| (Stay, choice) vs (Bad, fixation) | 2.52 (0.14) | 1.60 (0.16) | 9.20 | < .0001 | [0.72 to 1.12] | 0.92 |
| (Good, fixation) vs (Bad, fixation) | 1.59 (0.16) | 1.60 (0.16) | 0.00 | = .89 | [-0.27 to 0.24] | -0.02 |
| *Cluster 3* |  |  |  |  |  |  |
| Parametric bootstrap test (n = 10,000) | *p* |  |  |  |  |  |
| Full model vs. null model | < .001 |  |  |  |  |  |
| Post hoc<br>(pairwise t-test, Bonferroni correction) | Mean (SD) | Mean (SD) | *t* | *p* | 95% CI | Effect size |
| Contra |  |  |  |  |  |  |
| (Return, choice) vs (Stay, choice) | 2.97 (0.09) | 2.93 (0.09) | 1.08 | = .28 | [-0.04 to 0.14] | 0.05 |
| (Return, choice) vs (Good, fixation) | 2.97 (0.09) | 2.00 (0.11) | 12.31 | < .0001 | [0.82 to 1.13] | 0.98 |
| (Return, choice) vs (Bad, fixation) | 2.97 (0.09) | 2.02 (0.11) | 12.16 | < .0001 | [0.80 to 1.11] | 0.96 |
| (Stay, choice) vs (Good, fixation) | 2.93 (0.09) | 2.00 (0.11) | 11.38 | < .0001 | [0.77 to 1.09] | 0.93 |
| (Stay, choice) vs (Bad, fixation) | 2.93 (0.09) | 2.02 (0.11) | 11.22 | < .0001 | [0.75 to 1.07] | 0.91 |
| (Good, fixation) vs (Bad, fixation) | 2.00 (0.11) | 2.02 (0.11) | -0.20 | = .84 | [-0.22 to 0.18] | -0.02 |
| Ipsi |  |  |  |  |  |  |
| (Return, choice) vs (Stay, choice) | 2.83 (0.09) | 2.76 (0.09) | 1.48 | = .14 | [-0.02 to 0.16] | 0.07 |
| (Return, choice) vs (Good, fixation) | 2.83 (0.09) | 1.77 (0.12) | 12.07 | < .0001 | [0.89 to 1.23] | 1.06 |
| (Return, choice) vs (Bad, fixation) | 2.83 (0.09) | 1.78 (0.12) | 11.99 | < .0001 | [0.88 to 1.22] | 1.05 |
| (Stay, choice) vs (Good, fixation) | 2.76 (0.09) | 1.77 (0.12) | 11.15 | < .0001 | [0.82 to 1.17] | 0.99 |
| (Stay, choice) vs (Bad, fixation) | 2.76 (0.09) | 1.78 (0.12) | 11.06 | < .0001 | [0.81 to 1.16] | 0.98 |
| (Good, fixation) vs (Bad, fixation) | 1.77 (0.12) | 1.78 (0.12) | -0.11 | = .91 | [-0.24 to 0.21] | -0.01 |

## Discussion

In this study, several neurons within the anterior striatum demonstrated significant activity changed during the choice task, especially when subjects accepted good or rejected bad objects. By clustering the neurons according to their task-related responses, we observed that the activity of one cluster of neurons significantly increased while rejecting bad objects. The rejection strategies of the monkeys, including returning to a central point (return) and maintaining a gaze near it (stay), did not correlate with discernible differences in neural activity. This uniformity suggests that the neuronal processes underlying both rejection strategies are similar with respect to cognitive demand. Furthermore, the subdued neural activity observed during the fixation task relative to the choice task led us to posit that the neuronal activity associated with rejecting bad objects extends beyond mere reflexive saccade inhibition. This appears to be closely related to proactive inhibition, in which monkeys actively dismiss irrelevant options to achieve their goals.

### Involvement of cluster 2 neurons in cognitive control

Our findings revealed that neurons in cluster 2 exhibited heightened activity in response to bad objects, suggesting a pivotal role in their rejection. This response pattern aligns with the hypothesis that these neurons are integral to the indirect pathway, which is known to modulate inhibitory control. We observed no significant differences between the neural activities for the “return” and “stay” actions, indicating that these neurons were equally engaged for both rejection strategies.

Furthermore, the subdued neural activity of cluster 2 neurons during the fixation task underscores their specific involvement in proactive inhibition rather than mere suppression of reflexive saccades. This distinction highlights their role in evaluating and discarding suboptimal choices to facilitate goal-oriented actions, which is a cornerstone of adaptive behavior. These insights into the function of cluster 2 neurons underscore their contribution to the neural circuitry underlying proactive inhibition and cognitive control. Understanding the dynamics of these neurons sheds light on the mechanisms of cognitive regulation and has potential implications for addressing disorders characterized by impaired decision processes and inhibitory control.

### Anatomical and functional connections of task-related neurons

The task-related neurons identified in this study were predominantly located around the internal capsule of the anterior striatum, specifically in the ventral part of the head and body of the caudate nucleus and the dorsal part of the putamen. These regions receive extensive projections from the frontal eye field (FEF) and the supplementary eye field (SEF), which are critical areas for eye-movement control within the frontal cortex (Huerta et al., 1986; Stanton et al., 1988; Huerta et al., 1990; Shook et al., 1991; Parthasarathy et al., 1992). FEF and SEF neurons are implicated in the cancellation of actions during countermanding tasks, where a planned saccade should be suppressed (Hanes et al., 1998; Stuphorn et al., 2000), as well as for neurons involved in not selecting specific targets in the rostral FEF (Hasegawa et al., 2004). Furthermore, mild microstimulation of the SEF enhances performance during countermanding tasks by delaying saccade initiation (Stuphorn & Schall, 2006).

Given these connections, it is plausible that the neurons in cluster 2, which are associated with refusal of selection, may be influenced by inputs from these frontal cortex regions. This hypothesis is supported by a recent study that reported intensive modulation of caudate nucleus neuronal activity when monkeys canceled a planned saccade during a stop-signal task, which is analogous to a countermanding task (Ogasawara et al., 2018). These findings suggest that the striatal neurons in cluster 2 may play a crucial role during the proactive inhibition of actions and contribute to adaptive decisions based on the integration of sensory information and reward-based objectives.

### Role of the striatum during proactive and reactive inhibition

Previous human functional MRI studies have suggested that the striatum is involved in both proactive and reactive inhibition (Vink et al., 2005; Zandbelt & Vink, 2010; Pas et al., 2017). In contrast, the current study identified several neurons in the anterior striatum that were predominantly associated with proactive inhibition, with a notable absence of a response to reactive inhibition. However, this observation does not necessarily imply that the anterior striatum is not involved in reactive inhibition. Our experimental approach initially identified neurons that were responsive to the choice task and subsequently assessed their response to reactive inhibition during the fixation task. We may have overlooked neurons that are specifically responsive to reactive inhibition but not to proactive inhibition. This discrepancy raises the possibility that the striatum may be involved in reactive inhibition but operates through a mechanism different from that of proactive inhibition. Our results suggest a distinct neural basis for these two forms of inhibition within the striatum, suggesting that the cognitive processes underlying proactive inhibition and reflexive response suppression may be mediated by separate neural circuits (Jahanshahi et al., 2015).

### Integration of reactive and proactive inhibition in cognitive control

In our previous research, the neurons in the lateral part of the SNr demonstrated significant alterations in neural activity upon the rejection of bad objects during the choice task and suppression of saccades during the fixation task (Figures 9A and B; Yoshida & Hikosaka, 2023). Furthermore, local injections of glutamate receptor inhibitors into the lateral SNr, aimed at inhibiting excitatory projections from the STN, resulted in accelerated saccade responses to bad objects and frequent saccade suppression during the fixation task. These findings suggest the role of the STN-SNr pathway, a component of the hyperdirect pathway, in mediating reactive inhibition.

**Figure 9.**
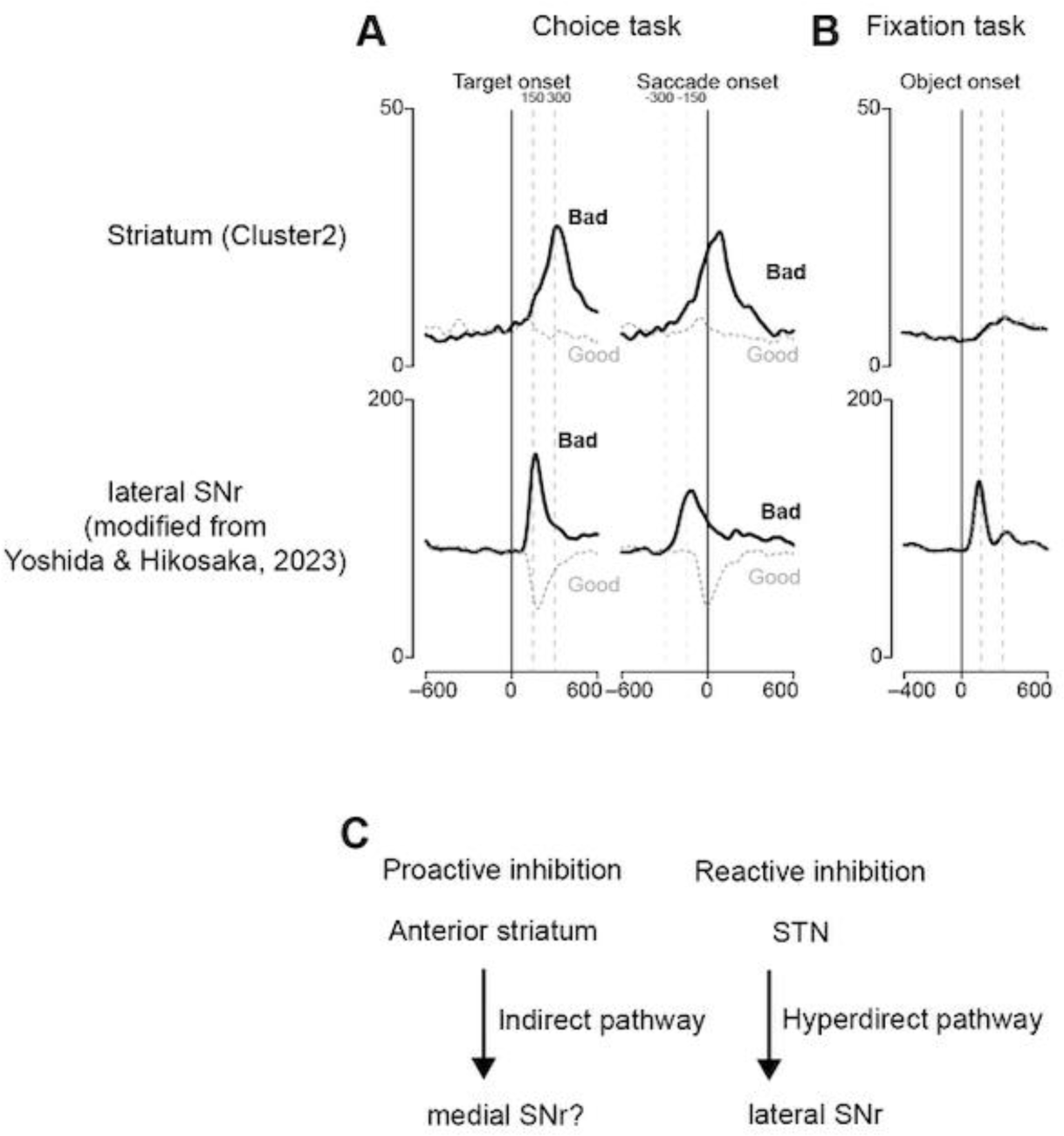
Comparison of putative inhibition-related striatal neurons with lateral SNr neurons. **(A)** Population activity of striatal (cluster 2) and SNr neurons aligned at the target or saccade onset during the choice task. **(B)** Population activity at object onset during the fixation task. **(C)** Hypothetical circuit diagram of two inhibitions.

In the present study, the neural activity of cluster 2 neurons, which was associated with the rejection of bad objects, peaked near the initiation of the return saccade. On comparing the neuronal activity associated with selective inhibition in the anterior striatum with that in the lateral SNr, we observed a marked difference in temporal dynamics. Specifically, the change in neuronal activity in the anterior striatum was slower than that in the lateral SNr. This distinction suggests that the hyperdirect pathway facilitates the early phase of inhibition, whereas the indirect pathway governs the later phase. This is consistent with the inhibition function hypothesis (Aron, 2011).

This discrepancy in the time course of neuronal activity between these two regions also suggests that the disinhibition of neuronal activity in the striatum is unlikely to directly generate the neuronal activity observed in the SNr. A possible explanation for this is the anatomical projection pattern within these regions. The task-related neurons identified in our study were primarily located in the anterior striatum, which is thought to project to the rostral part of the SNr (Smith and Parent, 1986; Szabo, 1970; Yasuda and Hikosaka, 2015). Based on this anatomical consideration, it is plausible that reactive inhibition involves the lateral SNr, whereas proactive inhibition may be more closely related to the rostral SNr (Figure 9C). This hypothesis is consistent with the distinct temporal profiles of neuronal activity observed in this study. Further investigations are warranted to elucidate the specific neural circuits involved in different types of inhibition.

### Conclusion

We identified a distinct group of neurons in the anterior striatum that demonstrated significant changes in neural activity during the rejection of a bad object as part of a choice task. These neurons demonstrated similar activity patterns when the object was rejected following and without a saccade, suggesting that their role extends beyond the mere suppression of saccadic movements. They are actively involved in the process of choice rejection, irrespective of the method of rejection. Furthermore, the minimal change in neuronal activity observed during the fixation task indicates that these neurons are not primarily engaged during reactive inhibition. Their activities are more consistent with proactive inhibition, which focuses on discarding unnecessary options to achieve a specific goal. These findings clarify the neural substrates of cognitive control, emphasizing the importance of the anterior striatum in orchestrating goal-directed behavior and highlighting the distinct mechanisms underlying different types of inhibition within the brain.

## Acknowledgments

This research was supported by the Intramural Research Program at the National Institutes of Health, National Eye Institute (1ZIA EY000415). MRI scanning was conducted in the Neurophysiology Imaging Facility Core (National Institute of Mental Health, National Institute of Neurological Disorders and Stroke, and National Eye Institute). We thank D. Parker, H. Warnock, G. Tansey, K. Allen-Worthington, A.M. Nichols, D. Yochelson, J. Fuller-Deets, and M. Robinson for technical assistance.

